# Synthetic genome expansion reveals a transcriptional constraint on genome size

**DOI:** 10.64898/2026.06.26.734810

**Authors:** Ning Lu, Xin Gao, Michael C. Lanz, Marianna E. Estrada, Shicong Xie, Gabriel E. Neurohr, Jan M. Skotheim

## Abstract

Genome sizes vary across eukaryotes, largely because of differences in non-coding DNA. However, the physiological effects of this excess DNA are unclear. We engineered budding yeast strains carrying up to 12.8 Mb of predominantly non-coding human DNA, doubling the genome without altering endogenous genes. Genome expansion slowed growth and increased cell size. Spike-in-normalized ChIP-seq and RNA-seq showed that added DNA recruited RNA polymerase II, diverting it from endogenous genes and lowering endogenous mRNA concentration. Ribosome profiling and proteomics revealed little translation from added sequences. A model linked this transcriptional competition to reduced ribosome activity and growth. Thus, excess DNA imposes a fitness cost by sequestering transcriptional resources, providing a mechanistic basis for selection against genome expansion in rapidly proliferating organisms such as yeasts and bacteria.

## Introduction

Eukaryotic genome sizes span several orders of magnitude yet show little relationship with organismal complexity. For example, why would an onion require five times more DNA than a human ^1^? This apparent discrepancy is known as the C-value paradox ^2,3^, where the C-value denotes the total DNA content of a haploid genome. Most genome-size variation arises from differences in non-coding DNA rather than gene number ^4,5^, raising the question of whether this DNA has functional significance or is simply genomic ‘junk’ ^6,7^. Some non-coding sequences regulate nuclear organization ^8^, global transcriptional capacity ^9^, and cellular physiology ^10^, and specific classes, such as enhancers ^11^, long non-coding RNAs ^12^, and structural RNAs ^13^, have well-established molecular functions. However, these represent only a small fraction of non-coding DNA, and the physiological consequences of the large variation in total non-coding DNA across eukaryotes remain unclear.

One explanation for genome-size variation is the nucleoskeletal theory, which proposes that DNA content supports nuclear volume independently of the information it encodes ^14^. This model is motivated by the striking linear correlation between DNA content and cell size across a roughly 100,000-fold range in eukaryotic cells ^15–18^. Because genome-size variation primarily reflects differences in non-coding DNA, such DNA has been proposed to act as structural material that scales nuclear volume with cell size. However, experimental manipulation of DNA content does not generally produce the predicted changes in relative nuclear volume. In fission yeast, for example, a 16-fold change in ploidy does not alter the nuclear-to-cell volume ratio ^19^. Together with models in which nuclear volume is determined primarily by nuclear protein content and osmotic balance ^20,21^, these findings argue against a predominantly structural role for excess DNA and suggest that genome expansion may influence physiology through other mechanisms.

To directly determine how non-coding DNA abundance affects cellular physiology, we developed a synthetic approach that expands the budding yeast (*S. cerevisiae*) genome without altering its endogenous gene content. We generated a series of strains carrying progressively increasing amounts of predominantly non-coding human DNA, adding more than 12 Mb and approximately doubling the native genome. This system allowed us to isolate the effects of non-coding DNA abundance from changes in gene content, ploidy, and chromosome number. We show that excess non-coding DNA slows growth, increases cell size, and delays the G1/S transition. The added DNA recruits RNA polymerase II and diverts it from endogenous genes, lowering endogenous mRNA concentration while contributing little to protein synthesis. Our results identify competition for transcriptional resources as a physiological cost of genome expansion and a selective force favoring compact genomes in fast-growing species.

## Results

Eukaryotic genomes exhibit extensive size variation, with larger genomes containing higher fractions of non-coding DNA ^22^. To examine how genome composition scales with genome size, we analyzed reported coding and total genome sizes from 153 previously published eukaryotic genomes ^23^ and calculated the corresponding non-coding DNA fractions. Consistent with previous reports, the fraction of non-coding DNA increased markedly with genome size, frequently exceeding 90% in large genomes, whereas compact genomes, such as those in yeasts, contained relatively little non-coding DNA (**Figure 1A**). Across unicellular eukaryotes, total DNA content correlates positively with cell size (**Figure 1B**), consistent with previous observations ^15–18^. These patterns raise the possibility that non-coding DNA contributes to fundamental cellular properties such as cell volume. However, in natural systems, variation in genome size is typically accompanied by concomitant changes in coding and regulatory sequences that make it difficult to isolate the causal effects of non-coding DNA (**Figure 1C**).

**Figure 1.**
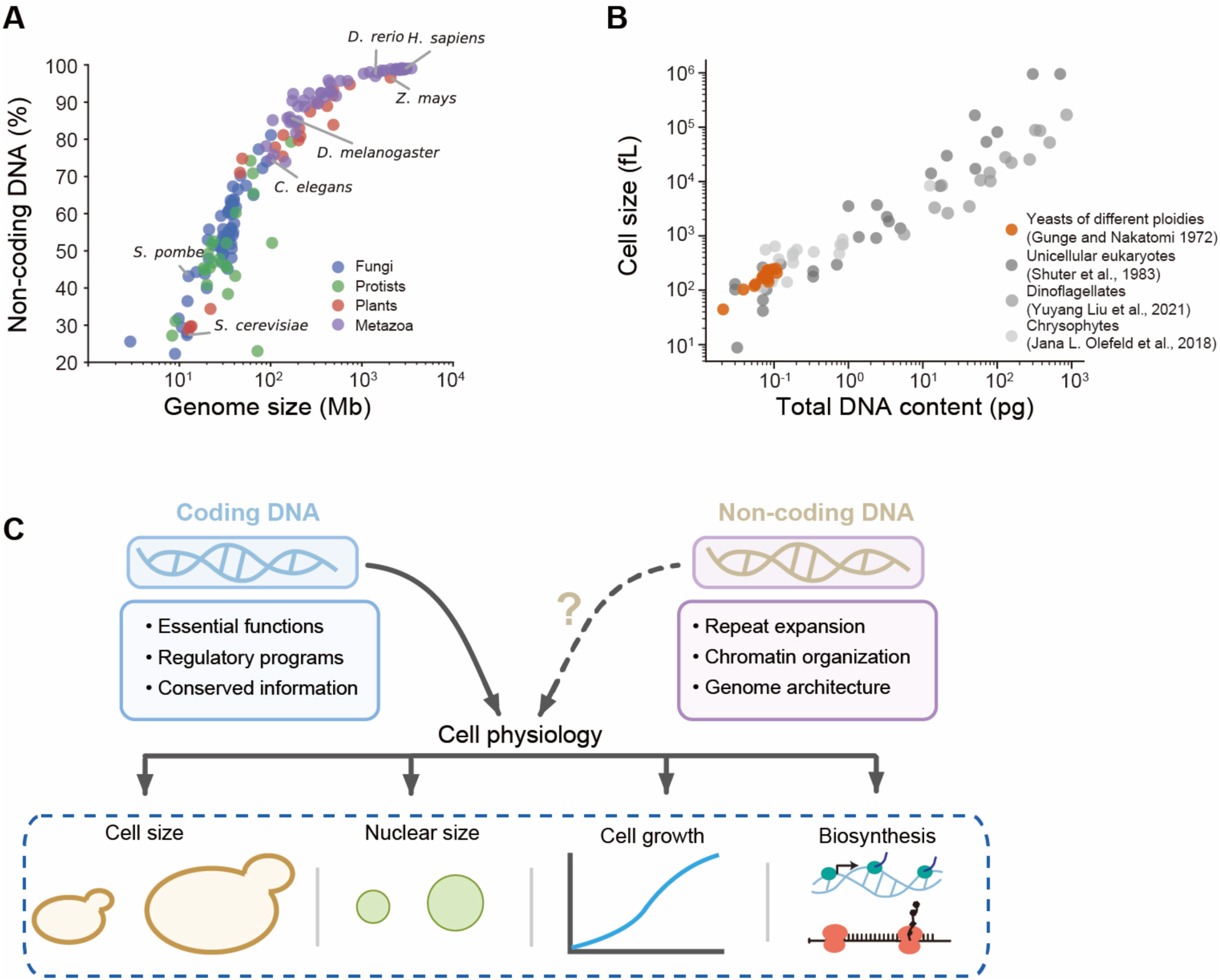
Potential links between genome size, non-coding DNA content, and cellular physiology across eukaryotes. **(A)** Scatter plot showing the fraction of non-coding DNA in 153 eukaryotic genomes ^23^. **(B)** Scatter plot showing genome size versus cell size for previously reported unicellular eukaryotes ^15–18^. **(C)** Schematic illustrating the possible impact of coding and non-coding DNA on cellular physiology.

### A synthetic platform for large-scale genome expansion

To directly examine the physiological consequences of non-coding DNA expansion, we built a synthetic platform to progressively increase genome size in *S. cerevisiae*. The platform is based on chromosomal fusion of YACs carrying large fragments of human DNA (∼0.85 Mb) derived from the Y chromosome (**Figures S1A-S1C**). Sequential YAC-chromosome fusions allowed stepwise insertion of large DNA fragments at different chromosomal termini. After fourteen rounds of integration, we generated strains containing up to ∼12.8 Mb of additional DNA, substantially increasing total genome content while preserving ploidy and endogenous gene composition (**Figure 2A**). During integration, the YAC centromere and telomeric sequences are simultaneously removed ^24,25^, allowing the host chromosome to support stable replication and segregation of the inserted DNA while maintaining the native chromosome number. Iterative integration was enabled by marker-assisted selection using additional amino-acid auxotrophic markers, allowing repeated YAC insertions while maintaining a single episomal YAC copy. To enable marker recycling after each round, a galactose-inducible Cre recombinase was introduced to excise loxP-flanked selection markers (**Figure 2B**).

**Figure 2.**
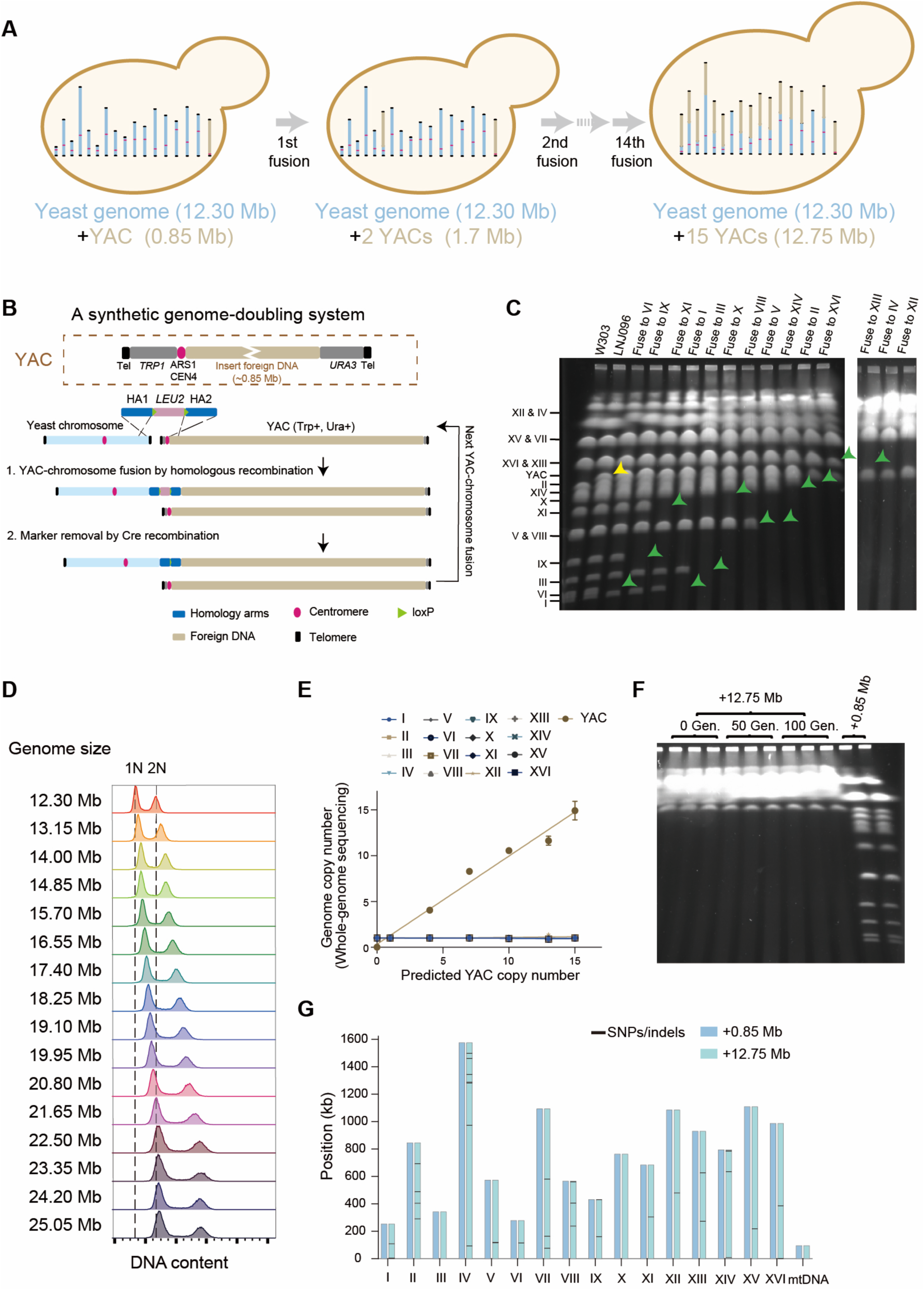
Generation of *S. cerevisiae* strains carrying increasing amounts of non-coding DNA. **(A)** Schematic illustrating the generation of a series of genome-expanded yeast strains through 14 rounds of sequential YAC-chromosome fusion. **(B)** Experimental schematic showing the YAC is duplicated and fused to one of the yeast chromosomes. In the first step, homologous arms flanking a *LEU2* selection marker are used to fuse the YAC to the targeted chromosome’s terminus while removing the telomeres. In the second step, the selection marker is excised using the Cre-loxP system, enabling subsequent rounds of YAC-chromosome fusion. **(C)** Pulsed-field gel electrophoresis showing bands with the indicated chromosomes. When a particular chromosome is fused to the YAC it gains ∼0.85 Mb and runs at a larger size. The yellow arrow indicates the position corresponding to the band size of the original YAC. Green arrows indicate where the targeted chromosome would have run if it were not fused to a YAC. Strains from left to right contain progressively more non-coding DNA following iterative YAC fusions. **(D)** The collection of YAC-containing yeast strains was stained with propidium iodide and examined using flow cytometry to measure DNA content. **(E)** Whole-genome sequencing-based measurements of chromosomal and YAC copy numbers. Data are shown as mean ± range (n = 2 biological replicates). **(F)** Strains carrying an additional 12.75 Mb of DNA were continuously passaged in SCD medium for 100 generations, and karyotype stability was assessed by PFGE. Three biological replicates are shown. **(G)** Distribution of mutations across yeast chromosomes, showing 27 single nucleotide polymorphisms (SNPs) and 15 small insertions or deletions (indels) identified in the final strains relative to the initial strain. Among these, a single SNP was detected within the coding sequence (CDS) of each of the following genes: *PMT2*, *LYS2*, *TDP1*, *RPB7*, *PDR15*, *THI74*, *CTA1*, *YFL015C* and *MUD2*, whereas all remaining mutations were located in non-coding regions of the yeast genome.

Successful chromosomal fusions were validated by pulsed-field gel electrophoresis (PFGE) after every fusion. When a particular chromosome is fused to the YAC it gains ∼0.85 Mb and runs at a larger size (**Figures 2C and S1D**). Diagnostic PCR verified the loss of native chromosome termini and the formation of YAC-chromosome junctions (**Figures S2A-S2C**). Genome expansion was assessed by flow cytometry, which revealed a progressive shift in DNA content distributions of fusion strains toward higher values, consistent with substantial increases in total genome content (**Figure 2D**). We further validated genome expansion by quantitative PCR and whole genome sequencing revealed that few mutations accumulated during strain construction (**Figures 2E, 2G, and S2D**). To evaluate the karyotype stability of the engineered strains, the strain carrying ∼12.8 Mb of non-coding DNA was passaged for over 100 generations. PFGE revealed no detectable changes in chromosome banding patterns between the original and passaged populations, indicating that the karyotype remained stable during prolonged propagation (**Figure 2F**).

### Genome expansion increases cell size primarily by delaying the G1/S transition

After establishing our strain series, we examined how non-coding DNA expansion affects cellular physiology. Cell volume increased nearly in proportion to total DNA content, so that the strains carrying twice the amount of total DNA as a wild type haploid were similar in size to diploid cells (**Figures 3A and S3A**). Moreover, an independent YAC accumulation system reported in the accompanying study ^26^ (Estrada et al. 2026, co-submitted paper) exhibited similar increases in cell size, indicating that the phenotype is unlikely to arise from sequence-specific properties of the introduced DNA. Taken together, these results suggest that increasing amounts of genomic DNA may drive an increase in cell size.

**Figure 3.**
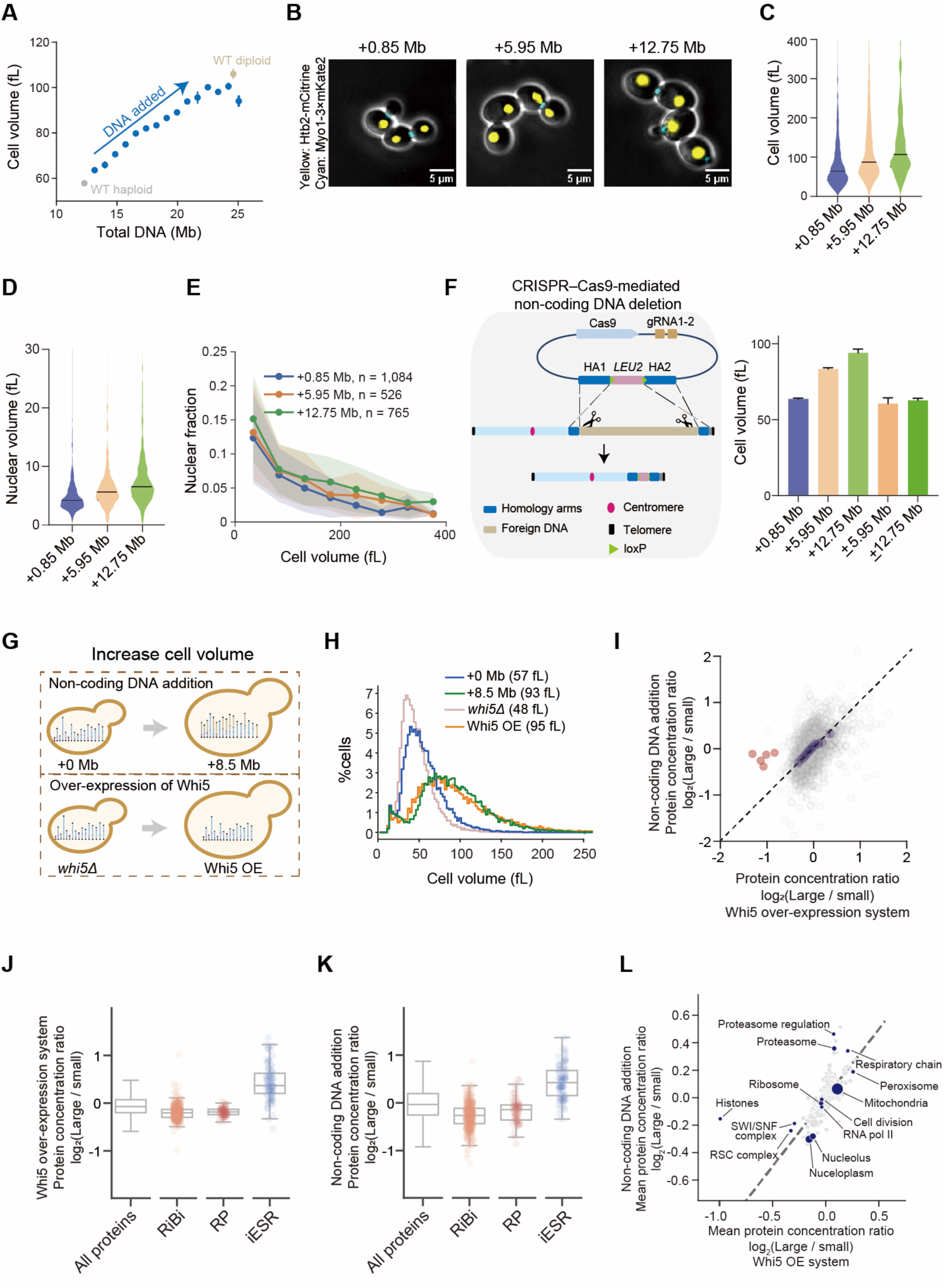
Non-coding DNA increases cell size. **(A)** Mean cell volume of asynchronously growing cultures in SCD medium measured by Coulter counter (n ≥ 2; mean ± s.d.). WT is a W303 strain. Blue dots correspond to strains containing increasing numbers of YACs. **(B)** Representative fluorescence microscopy images of cells with different non-coding DNA contents, showing Htb2-mCitrine (nuclear marker, yellow) and Myo1-3×mKate2 (bud neck marker, cyan). **(C)** Cell-size distributions measured by microscopy are shown as violin plots, with the median indicated by a central line. +0.85 Mb, n = 1,084; +5.95 Mb, n = 526; +12.75 Mb, n = 765. **(D)** Nuclear-size distributions measured by microscopy are shown as violin plots, with the median indicated by a central line. +0.85 Mb, n = 1,084; +5.95 Mb, n = 526; +12.75 Mb, n = 765. **(E)** Nuclear-to-cytoplasmic volume ratio as a function of cell volume determined by wide-field microscopy in SCD medium. Line plots indicate binned averages, shaded areas denote the mean with SD.| **(F)** Cas9 cleavage at both ends of the introduced DNA, followed by homologous recombination in yeast, enabled one-step removal of the non-coding DNA. Cell volume was measured before and after DNA removal. Data are shown as mean with range from at least two biologically independent replicates. **(G)** Schematic illustrating the cells whose proteomes will be compared. Cell size is increased by the addition of non-coding DNA or by the over-expression (OE) of the cell cycle inhibitor Whi5. **(H)** Distributions of cell size for the indicated genotypes measured by Coulter counter. Mean cell size is indicated in the legend. A representative histogram from two biological replicates is shown. **(I)** Correlation of protein concentration ratios derived from the indicated strains (n = 2 biological replicates per strain). Core histone proteins are highlighted in red. The identity line is shown as a dashed line. Blue points represent x-binned data, with error bars indicating the 99% confidence intervals. **(J)** Distribution of mean protein concentration ratios (n = 2) for genes associated with the rESR (repressed) and iESR (induced ESR) in the Whi5 over-expression system. The rESR comprises ribosomal protein (RP) and ribosome biogenesis (RiBi) gene clusters. ESR is the environmental stress response. **(K)** Distribution of mean protein concentration ratios (n = 2) for genes associated with the rESR and iESR in cells containing expanded non-coding DNA. **(L)** Annotation enrichment analysis of protein concentration changes (n ≥ 2 biological replicates) from cells that are larger due to either having more non-coding DNA (this study) or Whi5^29^. Each dot represents an annotation group; dot size reflects the number of proteins in the group, and position indicates the mean protein concentration change of the group.

We next asked whether genome expansion increased nuclear size beyond that expected from the accompanying increase in cell size, as proposed by the nucleoskeletal theory. To test this, we measured nuclear and cell sizes of histone labeled strains (*HTB2-mCitrine*) using live-cell microscopy. This showed that both nuclear and cellular volumes increased in genome-expanded strains (**Figures 3B-3D**). However, the nuclear-to-cytoplasmic ratio changed only modestly. When plotted as a function of cell volume, measurements from all strains fell along nearly the same curve, and cells of comparable size exhibited similar nuclear-to-cytoplasmic ratios regardless of genome size, consistent with previous experiments on fission yeast^19^ (**Figures 3E and S3B**). These results indicate that nuclear enlargement in genome-expanded strains is largely a side-effect of the increase in cell size, with little evidence that added non-coding DNA directly increases relative nuclear size. To test for the effect of any mutations arising during the expanded-genome strain construction, we removed the additional non-coding DNA from the ∼6 Mb and ∼12.8 Mb YAC strains using CRISPR-Cas9. This restored cell size to wild-type levels, demonstrating that these phenotypes arise directly from genome expansion (**Figures 3F and S3C**).

After finding that non-coding DNA has little effect on the nuclear-to-volume ratio, we next asked how it influenced other aspects of cellular composition. To measure cellular composition, we performed quantitative mass spectrometry and found widespread changes in diverse protein concentrations. We noted that these proteome changes were mostly similar to those we had found previously in larger cells ^27^. Since additional non-coding DNA increased cell size, we next sought to compare the proteomes of similar-sized genome-expanded cells and cells having only the endogenous genome. To do this, we controlled the expression of the G1/S cell cycle inhibitor *WHI5* ^28,29^ to increase the size of a haploid strain to match the size of a strain containing an additional 8.5 MB of YAC DNA (**Figures 3G and 3H**). We then compared the proteomes of these two large strains to that of normally sized cells. Strikingly, the overall proteomic changes observed in the two large cells were highly similar, indicating that non-coding DNA expansion produces global protein scaling effects comparable to those caused by cell size-mediated genome dilution (**Figures 3I-3L**). However, there are some notable differences. Genome-expanded cells have increased histones, likely generated in response to their additional DNA, and increased proteins associated with proteolysis, possibly in response to the generation of aberrant peptides from the non-coding DNA that need to be degraded.

After finding out that most of the proteome remodeling arose from the increased cell size of genome-expanded cells, we sought to identify the mechanism underlying this increase in cell size. To do this, we analyzed the DNA content distributions of our strain series using flow cytometry. Genome-expanded strains growing in SCGE media exhibited an increased proportion of cells in G1 phase relative to control strains (**Figure 4A**). To further test if non-coding DNA delayed the G1/S transition, we synchronized cells in G1 using α-factor and released them into the cell cycle. This revealed delayed progression through the G1/S transition after release, consistent with our flow cytometry data (**Figure 4B**). Time-lapse microscopy of individual cells further showed that genome expansion predominantly affected the relationship between cell size and cell cycle progression at the G1/S transition, whereas S/G2/M phases were only modestly affected (**Figures 4C-4E and S3D-S3G**).

**Figure 4.**
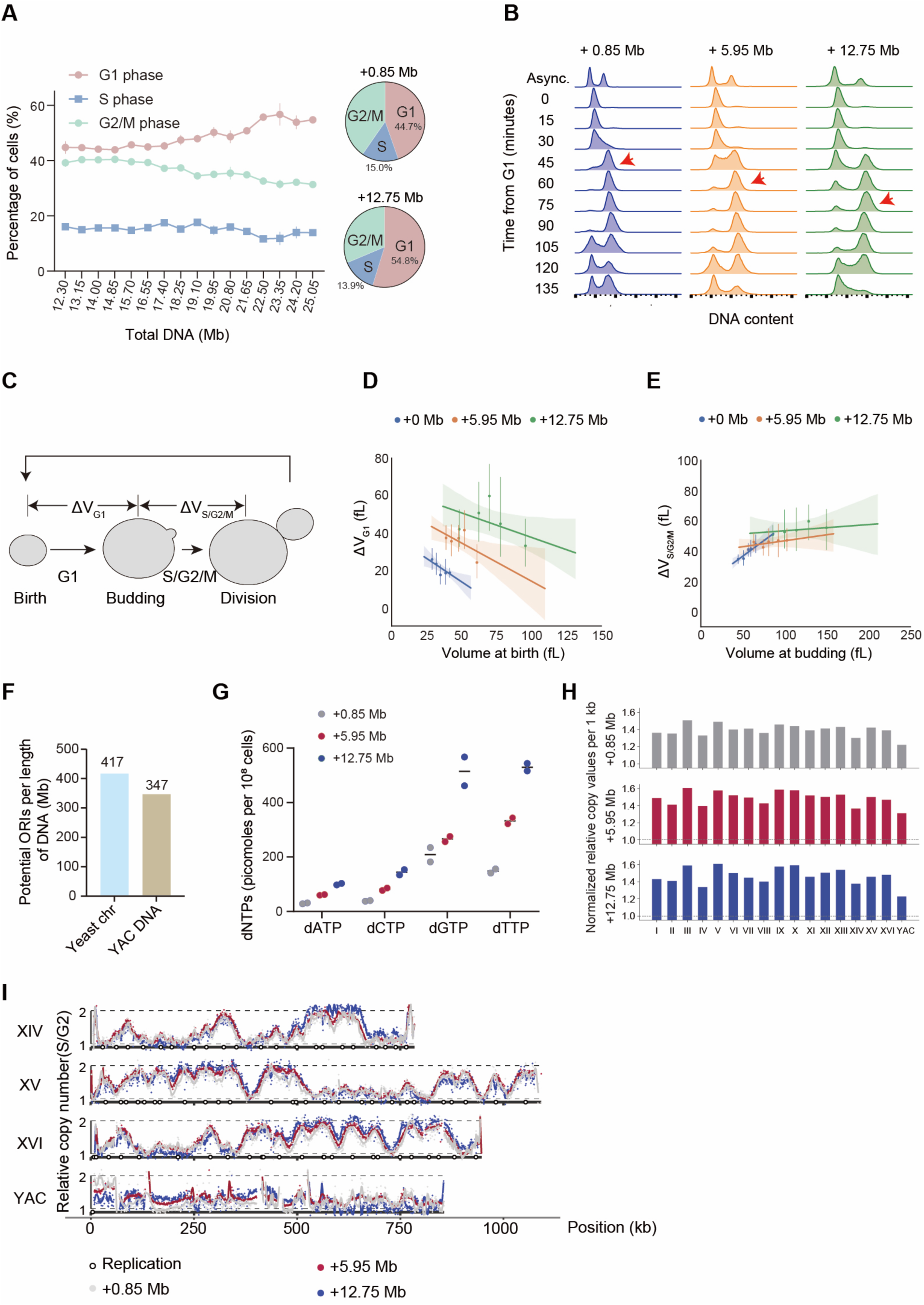
Non-coding DNA expansion delays G1/S transition without altering native chromosome replication dynamics. **(A)** Fraction of cells in G1, S, and G2/M phases as a function of non-coding DNA content determined by flow cytometry in SCGE medium and quantified using the Dean-Jett-Fox model in FlowJo. Data are shown as mean with range (n = 2). **(B)** Cell-cycle progression of α-factor-synchronized cultures analyzed by flow cytometry in SCGE medium. Samples were collected every 15 min, and a representative dataset from two biological replicates is shown. Red arrows indicate the time point cells fully transitioned from G1 to G2 phase following release. **(C)** Schematic of the first cell cycle of daughter cells analyzed by time-lapse microscopy at the single-cell level. **(D)** Single-cell growth dynamics of first-generation daughters profiled in SCGE medium using microfluidics. Volume growth from birth to budding plotted against birth volume. Data are binned by equal cell numbers; dots indicate mean per bin; error bars, 95% CI; solid line, regression to unbinned data; shaded area, 95% CI. +0 Mb, n = 80^48^; +5.95 Mb, n = 104; +12.75 Mb, n = 57. **(E)** Volume growth from budding to mitosis plotted against budding volume for the same cells as in **(D)**. **(F)** Predicted density of replication origins (ORIs) on the YAC and yeast chromosomes, determined using iORI-PseKNC2.0 ^34^. **(G)** Cellular dNTP levels quantified by liquid chromatography-mass spectrometry (LC-MS) in strains with increasing non-coding DNA content. Data are shown as mean with 2 biological replicates. **(H)** Sort-seq was performed using fluorescence-activated cell sorting (FACS) to enrich cells in S or G2 phases. FACS-enriched cells were collected, and genomic DNA was extracted from each cell population. Average replication differences across chromosomes and the YAC were compared by calculating normalized relative copy values per 1 kb. S/G2 scaling factors were applied to adjust read-count ratios to a 1-2 relative copy-number range. **(I)** Replication profiles of yeast strains carrying +0.85 Mb (grey points), +5.95 Mb (red points), and +12.75 Mb (blue points) of additional DNA are shown together with their respective spline-smoothed fits (grey, red, and blue lines). For direct comparison, replication profiles of *S. cerevisiae* chromosomes XIV, XV, and XVI and the YAC are overlaid across the three strains. Replication profiles of additional chromosomes are shown in **Figure S4**.

After identifying the G1/S transition as being delayed by non-coding DNA, we sought to test if any of the canonical G1/S regulators were responsible for this effect. To do this, we deleted either the G1 cyclin *CLN3*, a cell cycle activator, or *WHI5* from our set of genome-expanded strains. The set of *cln3Δ* strains was larger, and the set of *whi5Δ* strains was smaller, consistent with the effect of these mutations on cell size ^30–33^. In both these two new sets of strains, cell size still increases in proportion to the amount of added non-coding DNA indicating that the effect of non-coding DNA on the G1/S transition does not require either of these two key regulators (**Figures S3H and S3I**). Next, we performed time-lapse imaging of YAC-containing strains that lack *WHI5*. This showed a decrease in the amount of growth in G1 for a given cell size at birth and a more modest increase in S/G2/M growth (**Figures S3D-S3G)**. Taken together, these data provide further support that the G1/S transition is the primary conduit between non-coding DNA and increased cell size.

The modest extension we observed in S/G2/M, the anticipated effect of adding large amounts of extra DNA, and the prediction that YAC sequences have lower densities of replication origins^34^ prompted us to examine whether non-coding DNA affects S-phase progression (**Figure 4F**). However, intracellular dNTP pools scaled with total DNA content, suggesting upregulation of nucleotide biosynthesis and other DNA replication resources in response to the increased demand for them in genome-expanded strains (**Figure 4G**). Genome-wide replication timing analysis using Sort-Seq ^35^ revealed that early-firing origins on endogenous chromosomes did not exhibit noticeable decreases in replication peaks as YACs were added. Normalized relative copy values per 1 kb showed slightly higher values for endogenous chromosomes compared with YAC sequences, indicating modestly lower replication efficiency in the YAC regions (**Figures 4H, 4I, and S4)**. Thus, consistent with our single-cell analysis finding a lesser effect on S/G2/M phases of the cell cycle, we find that the integration of YAC fragments had minimal impact on the replication behavior of native yeast chromosomes. These observations, together with the accompanying study ^26^, argue against replication stress or DNA damage signaling as the primary cause of the cellular phenotypes associated with non-coding DNA accumulation. Taken together, these findings indicate that the introduction of large amounts of non-coding DNA is associated with a prolonged G1 phase and delayed G1/S transition, coinciding with an increase in cell size prior to S phase. This relationship reveals a quantitative link between genome content and cell cycle progression.

### Gene-expression output from excess non-coding DNA progressively declines from polymerase recruitment to peptide production

After establishing that genome expansion altered cell physiology, we next asked how the added non-coding DNA engages the gene-expression machinery and whether this engagement yields productive biosynthetic output (**Figure 5A**). Spike-in normalized ChIP-seq targeting the RNA polymerase II (RNAPII) subunit Rpb1 revealed widespread association of RNAPII with the integrated YAC sequences that was not restricted to the few coding regions of the human DNA (**Figures 5B and S5A**). As genome size increased, quantitative analysis showed that RNAPII occupancy (the global amount of DNA-bound RNAPII) on endogenous chromosomes per cell decreased substantially, whereas occupancy on YAC-derived sequences rose correspondingly (**Figure 5C**). These results indicate that excess non-coding DNA recruits RNAPII at a similar efficiency as yeast coding DNA to redistribute the limited pool of transcriptional machinery (**Figures 5D and S5B**). These findings are consistent with recent studies showing that random, synthetic, and evolutionarily naïve DNA sequences are broadly engaged by the yeast transcriptional machinery ^36–38^.

**Figure 5.**
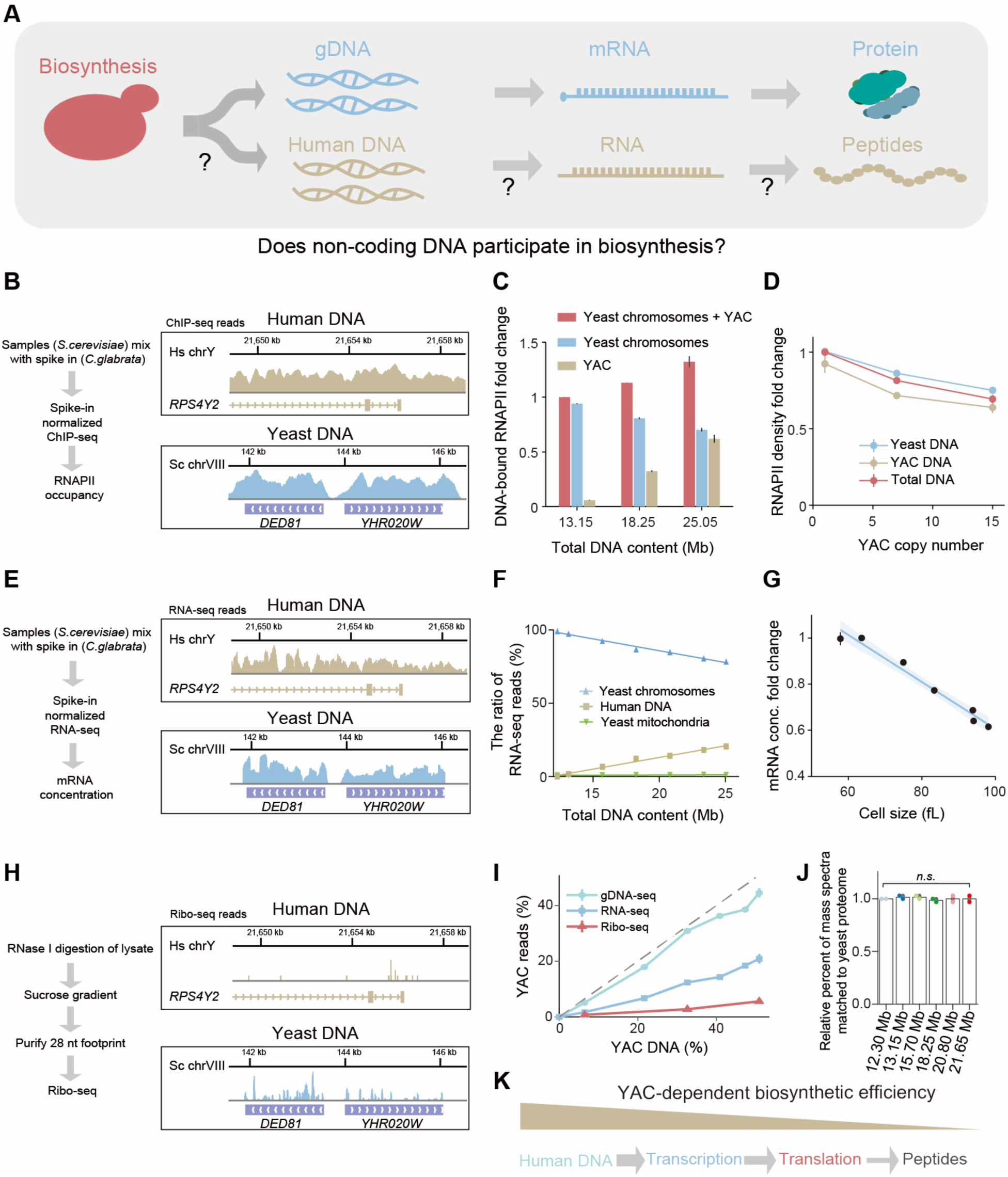
Expression of non-coding DNA progressively declines from RNA polymerase II recruitment to peptide synthesis. **(A)** Schematic illustrating the steps of expression of coding and non-coding DNA in yeast that will be measured using sequencing and proteomic methods. **(B)** Schematic of spike-in normalized ChIP-seq used to quantify global DNA-bound RNAPII in strains with increasing non-coding DNA. Representative Integrative Genomics Viewer (IGV) images showing the distribution of reads along the yeast genome and YAC DNA. **(C)** Bar plot showing total DNA-bound RNAPII per cell, including RNAPII associated with both the yeast genome and the YAC DNA. Data are shown as mean with range (n = 2). **(D)** Line plot showing the comparison of RNAPII density on yeast chromosomes versus YAC DNA. Data are shown as mean with range (n = 2). **(E)** Schematic of spike-in normalized RNA-seq used to quantify global mRNA levels in strains with increasing non-coding DNA. Representative IGV images showing the distribution of reads along the yeast genome and YAC DNA. **(F)** Fraction of RNA-seq reads mapping to yeast chromosomes, mitochondria, or the human Y chromosome as a function of non-coding DNA content (mean ± range, n = 2). **(G)** Total mRNA concentration per cell as a function of cell volume, determined by spike-in normalized RNA-seq (mean ± range, n = 2). The blue line shows a linear regression. **(H)** Workflow schematic of ribosome footprint sequencing. Representative IGV tracks showing the distribution of reads along the yeast genome and YAC DNA. **(I)** Line plots showing the fraction of reads mapping to the human Y chromosome from gDNA-seq, RNA-seq or ribosome profiling (Ribo-seq) as a function of non-coding DNA content. **(J)** Relative fraction of peptide spectral matches assigned to the yeast proteome across strains, determined by mass spectrometry. No increase in non-yeast peptides was detected. Two biological replicates were analyzed. **(K)** Schematic illustrating a progressive decrease in the expression of non-coding DNA.

To determine how the redistribution of RNAPII was reflected in RNA output, we performed spike-in normalized RNA-seq in strains carrying progressively more non-coding DNA. Reads mapping to YAC-derived sequences increased with genome size, whereas reads mapping to the endogenous yeast genome declined (**Figures 5E, 5F, and S5C**), consistent with the redistribution of RNAPII onto the YAC DNA. However, the fraction of RNAPII bound to YAC-derived sequences relative to genomic sequences was higher than the fraction of YAC-derived steady-state RNA relative to endogenous mRNA. This indicates that the inserted non-coding DNA recruits polymerase efficiently but supports comparatively inefficient production and accumulation of stable transcripts. Despite this shift, total mRNA per genome and total mRNA per cell remained largely stable as genome size expanded (**STAR Methods; Figures S5D-S5F**). Since the genome-expanded strains are larger, mRNA concentration decreased by ∼36% when genome size approximately doubled (**Figure 5G**).

We next asked whether transcripts produced from the added non-coding DNA contributed to protein synthesis. Ribosome profiling ^39^ revealed that only a small fraction of ribosome footprints mapped to YAC-derived transcripts (**Figures 5H, 5I, S6A, and S6B**). Moreover, these footprints were sparsely distributed across exonic, intronic, and intergenic regions rather than concentrated over canonical open reading frames. Consistent with this, our proteomic analysis could not detect any increase in the expression of proteins not encoded by the yeast genome (**Figure 5J**). Total cellular protein and RNAPII subunit concentration also remained largely unchanged across genome-expanded strains (**Figures S6C and S6D**).

Together, these findings show that excess non-coding DNA engages the gene-expression machinery but yields progressively declining output at each downstream step, from polymerase recruitment to RNA accumulation, ribosome engagement, and peptide production. Rather than increasing biosynthetic capacity, genome expansion acts primarily as a sink for transcriptional resources, diverting RNAPII away from endogenous genes while contributing little to productive protein synthesis (**Figure 5K**).

### Competition for RNA polymerase II between endogenous and non-coding DNA reduces transcription to constrain cellular growth

To test if the additional non-coding DNA places a burden on the cell, we measured growth rates. We found that increasing amounts of non-coding DNA proportionally reduce the cellular growth rate (**Figure 6A**). To explain this, we developed a quantitative model of RNAPII binding and mRNA production in which endogenous yeast DNA and inserted YAC DNA compete for a shared pool of polymerase (**Figures 6B and 6C**). This model was motivated by the proportional decrease in total yeast mRNA concentration with increasing non-coding DNA (**Figure 5G**).

**Figure 6.**
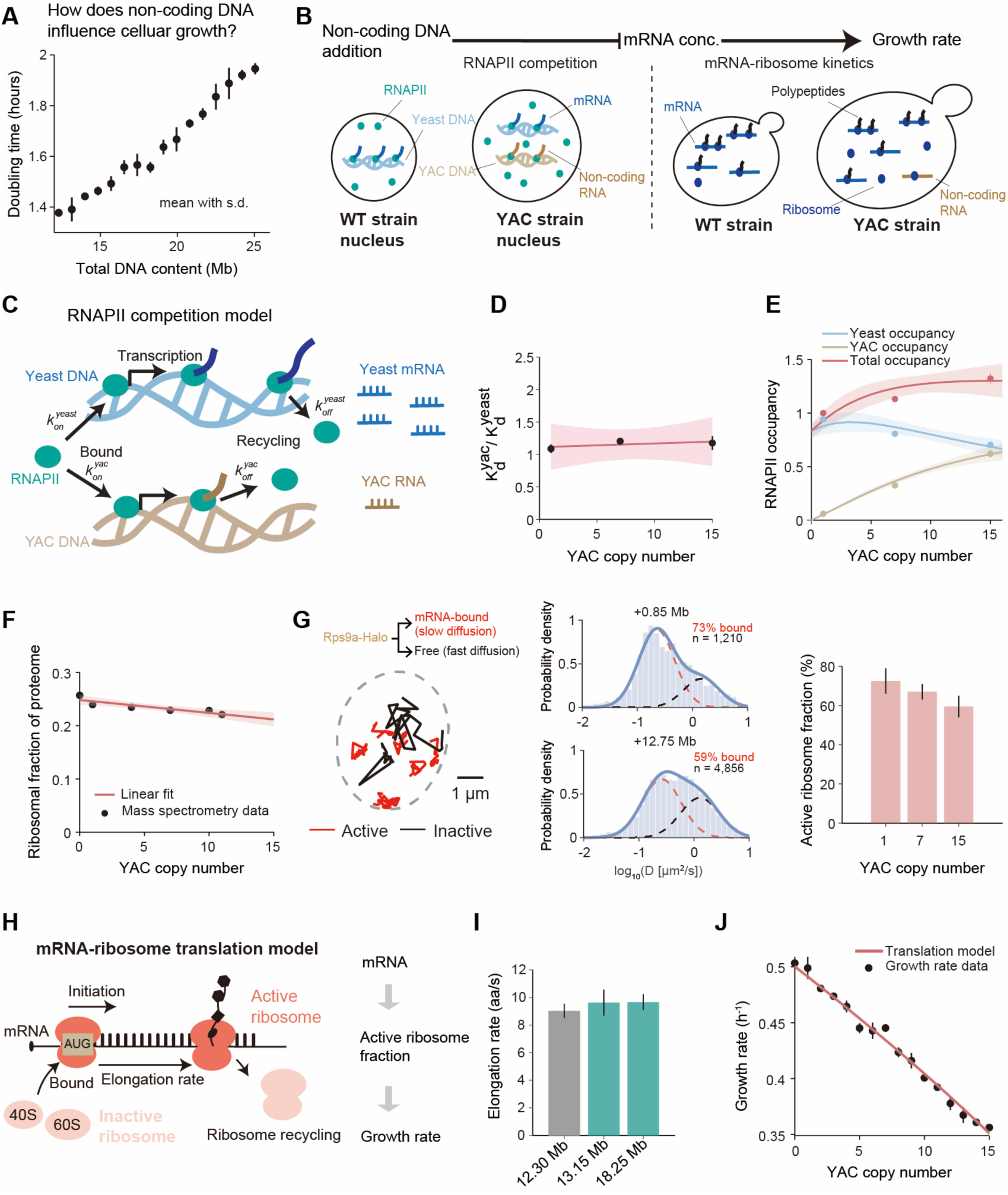
Competition with non-coding DNA for RNAPII leads to decreased transcription and cellular growth. (A) Doubling time determined from OD_600_ growth curves (n ≥ 2) using linear regression fits. (B) Schematic illustrating the competition between yeast and YAC DNA for RNAPII, and the mRNA-ribosome mass action kinetic model for translation. (C) Schematic illustration of competition for RNAPII by the yeast genome and YAC DNA. (D) Ratio of RNAPII dissociation constants between YAC and the yeast genome in the specified strain. Data are mean ± range (n = 2). The shaded band indicates the 90% CI of the linear fit. (E) DNA-bound RNAPII on yeast DNA, YAC DNA, and total DNA was fit with our RNAPII competition model. The shaded band indicates the 90% CI of the fit. (F) Ribosomal fraction of the proteome in the indicated YAC-containing strains (mean with range, n = 2). The shaded area denotes the 90% CI of linear fit. (G) Representative trajectories of DNA-bound (red) and unbound (black) ribosomes measured by single-molecule tracking, with the DNA-bound population used to calculate the active ribosome fraction (right hand panel). Actively translating ribosomes likely diffuse significantly slower than free ribosomes because of the relatively large size of mRNA. The fractions of active and inactive ribosomes were distinguished using a Gaussian mixture model to fit the distributions of diffusion coefficients measured from single-molecule tracking experiments. Distribution of diffusion coefficients of ribosomes in strains with 1 YAC (n = 1,210 tracks) and 15 YACs (n = 4,856 tracks), respectively. (H) Schematic of the detailed mRNA-ribosome translation model. (I) Comparison of elongation rates (aa/s) across strains with different amounts of non-coding DNA, calculated from experimental data (**see STAR Methods**). Data are shown as mean and s.e.m. (J) Cellular growth rate (ln(2)/population doubling time) of strains with the indicated number of YAC copies was fit using the mRNA-ribosome kinetic model based on mRNA and ribosome concentrations^40^. The shaded area denotes 90% CI of the fit.

In our model, the amounts of both yeast DNA-bound RNAPII 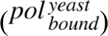 and YAC DNA-bound RNAPII 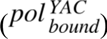, are determined by the mass action kinetics of free RNAPII, *pol_free_*, and the yeast and YAC DNA respectively. Dissociation of DNA-bound RNAPII is modeled using first-order kinetics, with the off-rate 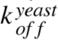 for yeast DNA-bound RNAPII and 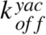 for YAC DNA-bound RNAPII. At steady state, the amount of both yeast and YAC DNA-bound RNAPII should follow:

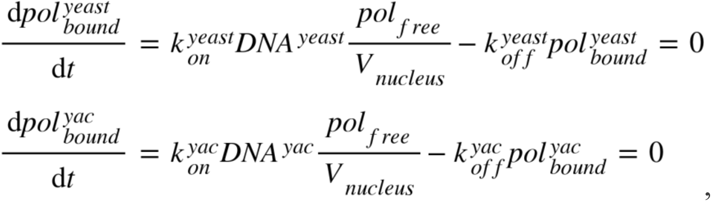

Where *V_nucleus_* denotes the nuclear volume, and 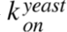 and 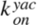 denote the binding rate of RNAPII to yeast and YAC DNA, respectively. Based on the observation that RNAPII concentration *c* remains nearly constant upon DNA addition (**Figure S6E**), the total RNAPII amount should be proportional to cell size *V_cell_*, so that:

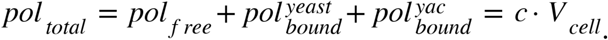

These three equations can be solved for the amount of endogenous yeast DNA-bound and YAC DNA-bound RNAPII, respectively, so that:

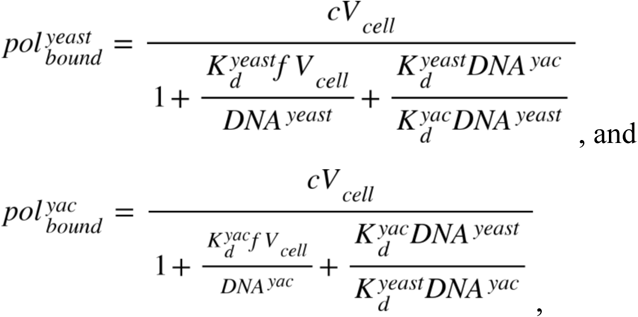

where the equilibrium dissociation constant of RNAPII with yeast DNA is denoted as 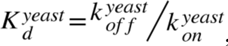, and that with YAC DNA is denoted as 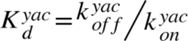. We note that the ratio of equilibrium dissociation constants of RNAPII on yeast DNA and YAC DNA is

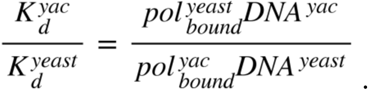

We can calculate this ratio from our RNAPII occupancy measurements from spike-in ChIP-seq experiments. We found that the equilibrium dissociation constant of RNAPII for YAC DNA is consistently ∼1.15-fold higher than that for yeast DNA across strains with different amounts of YAC DNA (**Figure 6D**). This indicates that RNAPII binds yeast DNA with a slightly stronger affinity than YAC DNA, as might be expected since these sequences have evolved to support specific transcription. Next, we measured cell size, nuclear fraction, and DNA content (**Figure S7A**), which greatly constrained our model and left only two free parameters, 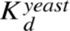, and the total RNAPII concentration *c* to fit the RNAPII occupancy on both yeast and YAC chromosomes. The fitted model accurately recapitulated the observed redistribution of RNAPII, with occupancy progressively increasing on YAC DNA and decreasing on endogenous chromosomes as genome size expanded (**Figure 6E**). These results support the idea that added non-coding DNA competes directly with the endogenous genome for RNAPII, thereby reducing transcriptional output from native genes.

Because steady-state mRNA levels reflect both synthesis and degradation, we next tested whether mRNA decay rates were affected by genome expansion. We repressed transcription of the methionine-regulated genes *MET3* and *MET17* and quantified mRNA decay by RT-qPCR. Decay rates were reduced in genome-expanded strains (**Figures S7B-S7D)**, consistent with previous observations that mRNA turnover slows in enlarged cells ^28^. Thus, the decrease in endogenous mRNA concentration cannot be explained by accelerated mRNA decay and instead arises despite partial compensatory stabilization of transcripts.

We next asked how reduced transcriptional output could influence growth. Although ribosome concentration has classically been viewed as a major determinant of growth rate, ribosome concentration in genome-expanded strains decreased only slightly, as measured by quantitative proteomics (**Figure 6F**). This small reduction in the ribosome concentration is insufficient to explain the ∼30% reduction in growth rate when ∼12.8 Mb of non-coding DNA was added. Since our previously established eukaryotic growth framework posits that the growth rate depends on both mRNA and ribosome concentrations ^40^, we sought to test if the decrease in total mRNA concentration could sufficiently reduce the fraction of actively translating ribosomes to sufficiently reduce the growth rate. We therefore quantified the fraction of actively translating ribosomes by single-molecule tracking using our previously described methods ^40^ (**Figures 6G and S7E**). The proportion of active ribosomes progressively decreased with increasing non-coding DNA content, indicating that translational capacity is not primarily limited by ribosome abundance, but by the reduced availability of mRNA in strains containing more non-coding DNA. Our eukaryotic growth model ^40^ requires the peptide elongation rate of single translating ribosomes, which could in principle be affected by non-coding DNA. To test this, we estimated the peptide elongation rate as described previously ^40^. We used single-molecule imaging to determine the fraction of ribosomes that were active so that our mass spectrometry data could be used to estimate the total number of translating ribosomes. The total protein synthesis rate is determined by the cellular growth rate, which allows us to estimate the protein synthesis rate per active ribosome. We find that the peptide elongation rate is ∼9.5 aa/s, consistent with previous results for wild type strains ^40^, and independent of the amount of non-coding DNA. Incorporating our measured peptide elongation rates (**Figures 6H and 6I**) and refitting only a single parameter corresponding to the mRNA concentration in a wild type strain, the model precisely reproduced the observed decrease in growth rate across genome-expanded strains (**Figure 6J**). As an independent test of the model, we reduced global transcription in non-genome-expanded yeast with thiolutin, an inhibitor of RNA polymerase ^41^. RNA fluorescence in situ hybridization confirmed reduced total mRNA levels, and this perturbation decreased growth rate while increasing cell size, thereby phenocopying the effects of genome expansion (**Figures S7F-S7K)**.

Together, these results support a model in which excess non-coding DNA competes with endogenous chromosomes for RNAPII, reducing native transcriptional output and lowering the fraction of active ribosomes. In this way, the reduction of productive gene expression explains the effect of non-coding DNA on cellular growth (**Figure 7**).

**Figure 7.**
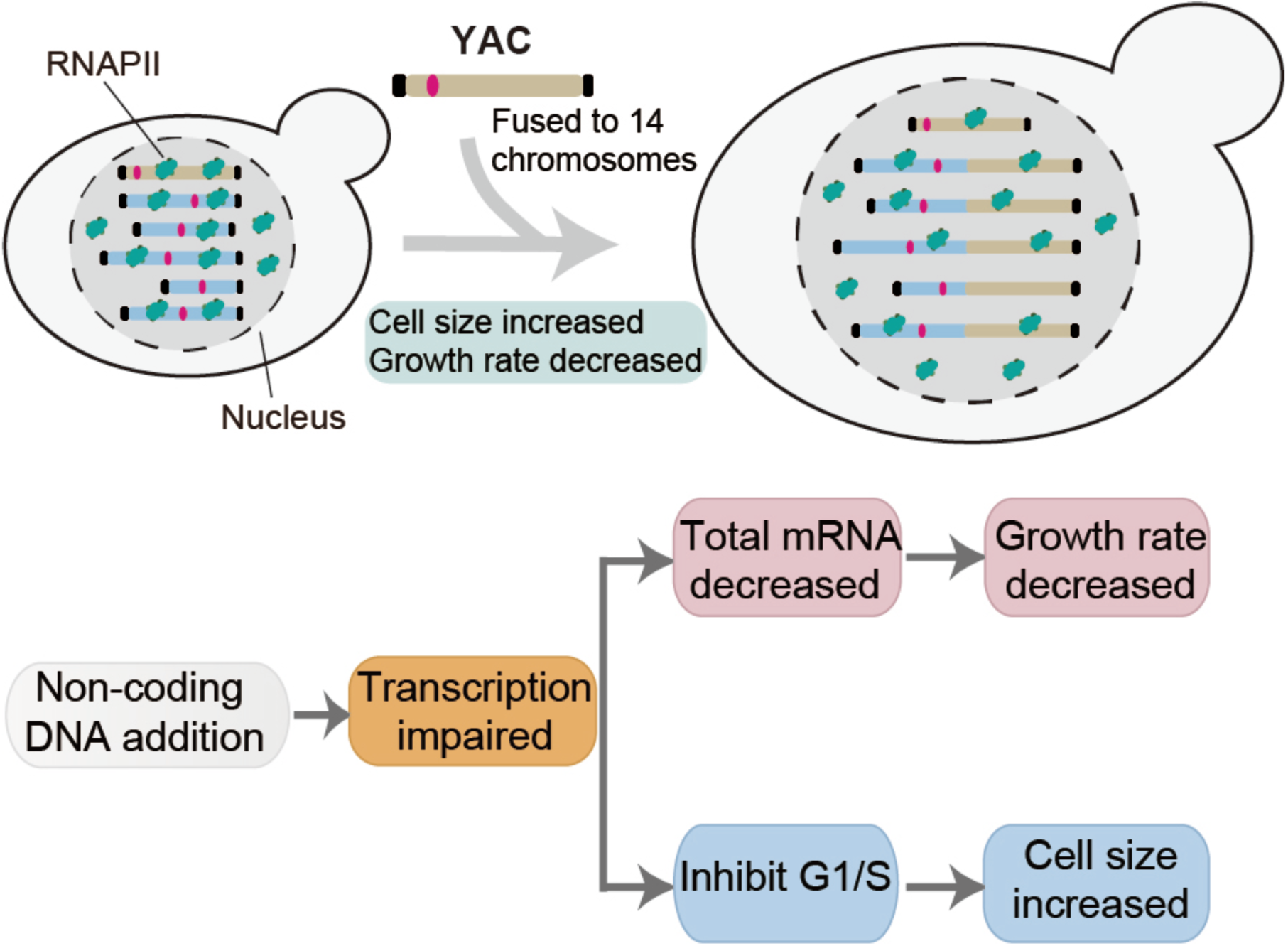
Increased non-coding DNA content impairs transcription and drives reduced growth with increased cell size. Model illustrating the physiological consequences of genome expansion through sequential YAC-chromosome fusions. Fourteen successive fusions introduced up to ∼12.8 Mb of additional non-coding DNA at chromosome termini while preserving ploidy and endogenous gene composition. The added DNA recruits RNA polymerase II (RNAPII), reducing its availability at endogenous genes and thereby lowering endogenous mRNA concentration. Reduced mRNA concentration slows growth, whereas delayed G1/S progression leads to increased cell size.

## Discussion

Our results show how the expansion of non-coding DNA can quantitatively influence cellular physiology by redistributing limited transcriptional resources. Although pervasive transcription of non-coding DNA was previously observed ^42–44^, its functional consequences were unclear. By developing a synthetic platform to expand non-coding DNA in budding yeast while maintaining the endogenous genome, we were able to isolate the effects of non-coding DNA from confounding variables such as gene content, ploidy, and chromosome number ^45–47^.

Expansion of non-coding DNA in budding yeast increased cell size, consistent with cross-species correlations of genome content and cell volume observed across unicellular eukaryotes ^15–17^. Spike-in normalized ChIP-seq and RNA-seq revealed that RNA polymerase II redistributes from the endogenous genome to YAC-derived non-coding sequences. This redistribution resulted in a ∼36% decrease in total mRNA concentration when the amount of YAC sequence was similar to the endogenous budding yeast genome. Ribosome profiling and mass spectrometry further showed that transcripts derived from the YAC DNA contribute minimally to protein synthesis, indicating that excess non-coding DNA acts primarily as a transcriptional sink. Mathematical modeling further supported that this transcriptional competition mechanism constrains growth in genome-expanded strains. By decoupling coding and non-coding DNA contributions, our platform enables rigorous causal inference that is not achieved by comparing diverse eukaryotic cells.

An important question is how broadly this mechanism applies across eukaryotes with much larger amounts of non-coding DNA. Our results do not imply that all large-genome organisms experience the same degree of transcriptional limitation as budding yeast. Rather, the burden imposed by excess DNA is likely to depend on how effectively it is mitigated. In many eukaryotes, non-coding sequences may be more efficiently silenced through chromatin-based repression, reducing their access to RNA polymerase II and limiting unproductive transcription. In addition, larger genomes are often associated with larger cells, which can accommodate greater absolute amounts of transcriptional machinery so that enough remains to support useful mRNA production. Both features would be expected to mitigate the impact of excess DNA on gene-expression capacity. However, neither buffering strategy is free: silencing must be actively maintained, and increasing transcriptional capacity requires additional biosynthetic investment. Thus, rather than representing a yeast-specific phenomenon, our findings point to a broader connection between genome expansion and the cellular resources required to repress or service excess DNA that are likely supported by larger cell sizes.

Finally, our findings suggest a mechanistic explanation for why the fastest-growing cells carry so little non-coding DNA. In organisms such as yeast and bacteria, where rapid proliferation depends on maximizing biosynthetic efficiency, excess DNA that engages transcriptional machinery without contributing meaningfully to protein production would be expected to reduce fitness. From this perspective, compact genomes may be favored not only because they are faster to replicate, but also because they minimize the diversion of transcriptional resources into non-productive outputs.

## STAR★METHODS

### Strain construction and yeast media

Our strains were based on AA17392 **(Figure S1C**). YACs contain the regions of the human Y chromosome as described in Foote et al ^49^. Genotypes of all strains used in this study are provided in **Table S1**. *S. cerevisiae* strains were generated using a standard lithium acetate transformation method. LNJ096 (+0.85 Mb) was generated by homologous recombination using a fusion PCR fragment containing a galactose-inducible promoter driving the Cre recombinase gene. This was integrated into chromosome XVI at the intergenic region between YPL160W and YPL159C. After integration, the *LEU2* auxotrophic marker was excised by expressing the galactose-induced Cre recombinase. Subsequent YAC-chromosome fusion strains were generated by transformation with fusion PCR fragments carrying homology arms to enable homologous recombination to fuse the YAC end to chromosomal termini. During fusion, the YAC centromere and a telomere from the YAC and the targeted chromosome were removed to stabilize the fused chromosome. In parallel, selective conditions (SCD medium lacking tryptophan and leucine) were applied to select for the maintenance of an additional free copy of the YAC. The additional free YAC can then be used for iterative fusions to create our strain series containing stepwise increases in the amount of exogenous DNA.

Cells were grown in synthetic complete medium (SC) supplemented with either 2% glucose (SCD) or 2% glycerol plus 1% ethanol (SCGE) as the carbon source. Unless otherwise specified, cells were cultured in SC supplemented with 2% glucose. For DNA content analysis (**Figure 2D**), cell-cycle analysis and single-cell dynamics (**Figures 4A, 4D, 4E, and S3D-S3G**), SC supplemented with 2% glycerol and 1% ethanol was used, as these conditions increase the range of cell sizes and the proportion of G1-phase cells subject to cell size control. All other experiments were conducted in SC supplemented with 2% glucose. Cells were maintained at 30 °C, cultured below an OD_600_ of 0.45, and harvested at OD_600_ values between 0.1 and 0.4.

### Karyotype analysis by pulsed-field gel electrophoresis (PFGE)

Karyotype analysis by PFGE was adapted from Shao et al ^50^. Yeast strains carrying YAC-chromosome fusions were cultured in 50 ml SCD medium at 30 °C until log phase. Cells were then harvested, washed sequentially with distilled water and 50 mM EDTA (pH 8.0), washed once with resuspension buffer (10 mM Tris-HCl, pH 7.2), and finally resuspended in 150 µl of the same buffer. The suspension was mixed with 75 µl Zymolyase-20T solution (20 mg/ml Zymolyase-20T (MP Biomedicals), 2.5% glucose, 50% glycerol, 50 mM Tris-HCl, pH 8.0), followed by addition of 225 µl 2% low-melting-point agarose dissolved in TE25S buffer (25 mM Tris–HCl, pH 8.0, 10.3% sucrose, 25 mM EDTA, pH 8.0). The mixture was cast into plug molds and solidified at 4 °C for 30 min. The plugs were then removed from the molds and incubated in 5 ml lyticase buffer (50 mM EDTA, pH 8.0, 10 mM Tris-HCl, pH 7.5) containing 500 µl Zymolyase-20T solution at 37 °C for 3 h. Next, the plugs were washed sequentially with distilled water and wash buffer (20 mM Tris-HCl, pH 8.0, 50 mM EDTA, pH 8.0) before being treated with proteinase K solution (100 mM EDTA, pH 8.0, 1% sodium lauryl sarcosine, 0.2% sodium deoxycholate, 1 mg/ml proteinase K) at 50 °C for 36 h. Finally, the plugs were washed four times with wash buffer at room temperature. Each wash lasted 1 h with gentle agitation. Chromosomal DNA was separated by PFGE on a 1% agarose gel (Bio-Rad) with a switch time of 60 s for 28 h, followed by 90 s for 14.5 h, at 6 V cm⁻¹ and 14 °C, and gels were subsequently stained with MilliporeSigma™ GelRed™ Nucleic Acid Stain (Catalog No. SCT123) to visualize chromosomal DNA bands.

### Quantification of YAC terminal copy number by quantitative PCR (qPCR)

Genomic DNA was extracted from yeast cells using a commercial kit (YeaStar Genomic DNA Kit; Zymo Research) according to the manufacturer’s instructions. Quantitative PCR was performed with iTaq Universal SYBR Green Supermix (Bio-Rad) using two primer pairs: one targeting the *ACT1* locus as an internal reference and the other targeting the terminal region of the YAC. Cycle threshold (Ct) values, defined as the PCR cycle at which fluorescence crossed a manually set threshold line within the exponential phase of amplification, were obtained for each primer pair. The relative copy number of the YAC terminal was calculated using the ΔCt method, where ΔCt = Ct (YAC terminal) – Ct (*ACT1*), and relative copy number = 2^-ΔCt^.

### Cell size measurements

Cells were grown to log phase (OD_600_ = 0.1-0.2), and 1 ml of culture was transferred to an Eppendorf tube and placed on ice. The cells were fully dispersed by sonication for 8-10 s. Subsequently, 100-200 μl of sonicated suspension was diluted in 10 ml Isoton II diluent (Beckman Coulter) prior to measurement. Cell size was determined using a Beckman Coulter Z2 particle counter. Two measurements (10,000-30,000 particles each) were averaged per sample. Particles < 10 fL or > 600 fL were excluded.

### Growth rate measurement

Single colonies were inoculated into 30 ml SCD medium and cultured at 30 °C. Cultures were maintained in exponential phase by continuous dilution, with measurements initiated at OD₆₀₀ ∼0.02. OD₆₀₀ was recorded hourly. Specific growth rates were calculated from the slope of a linear fit to log-transformed OD₆₀₀ values over time, and doubling times were derived as ln(2)/growth rate. At least two biological replicates were performed per strain.

### Measurement of the nuclear-to-cell volume fraction by microscopy

Yeast strains expressing Htb2-mCitrine and Myo1-3×mKate2 proteins were grown in SCD at 30 °C until reaching log phase (OD_600_ = 0.1-0.2). Cells were collected by centrifugation, gently resuspended, and placed onto 1% agarose pads (ibidi).

Images were acquired using a Zeiss Axio Observer.Z1 wide-field epifluorescence microscope fitted with a Colibri LED module and a 63×/1.4 NA oil-immersion objective. Htb2-mCitrine was excited with the 505 nm LED (25% intensity, 400 ms exposure), and Myo1-3×mKate2 with the 555 nm LED (25% intensity, 1 s exposure). Phase-contrast imaging was used to define cell boundaries. Temperature was maintained at 30 °C during acquisition using a TempController 37-2 unit. Z-stacks were acquired and the best-focused plane was manually selected. More than 500 cells were analyzed per replicate.

Cell segmentation was carried out automatically using ACDC software ^51^ and refined manually. Nuclear regions were identified from fluorescence images using a custom Python-based pipeline. Cell volume was estimated by approximating each yeast cell as a prolate spheroid, with cross-sectional areas perpendicular to the long axis integrated along its length. For budding cells, the mother and bud volumes were calculated independently and subsequently combined. Nuclear volume was estimated by approximating the nucleus as a tri-axial ellipsoid: the long (a) and short (b) axes were measured in two dimensions using the skimage package, while the third axis (c) was defined as *c* = (*a* + *b*) / 2, and nuclear volume was calculated as 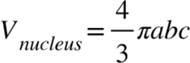. The nuclear-to-cell volume fraction was determined by dividing the nuclear volume by the cell volume.

### Live cell microscopy and image analysis

Cells were grown in SCGE medium to log phase (OD_600_ = 0.1-0.3), briefly sonicated, and introduced into a CellASIC Y04C microfluidics plate (Millipore Sigma) with continuous medium flow maintained at 2 psi^52^. Unless stated otherwise, time-lapse images were acquired every 6 or 15 min on an Observer Z1 (Zeiss) with an automated stage, an Axiocam 705 mono camera and a Plan-Apochromat 63×/1.4 NA oil-immersion objective at 30 °C. Myo1-3×mKate2 was used as a marker of budding onset, which coincides with the initiation of DNA replication^53^. Images of this marker were acquired with 500 ms or 200 ms exposures using a Colibri 555 nm LED module at 16% power. Segmentation and lineage tracking were performed with the Cell-ACDC analysis framework^51^. Mother-bud pairs were assigned within Cell-ACDC, and the timing of budding and cytokinesis was determined from the appearance and disappearance of Myo1-3×mKate2, respectively.

### Cell cycle progression assay

Yeast strains were grown in synthetic complete medium supplemented with 2% glycerol and 1% ethanol, and synchronized in G1 phase by the addition of α-factor to a final concentration of 5 µg/ml. A second α-factor treatment (5 µg/ml) was applied after 2.5 h, and cultures were harvested 2.5 h later by vacuum filtration to remove the arresting agent.

Cells were resuspended in pre-warmed fresh SCGE medium incubated at 30 °C. Aliquots (400 µl) were collected every 15 min, and fixed by adding 1 ml ice-cold 100% ethanol at 4 °C for ≥1 h. Cells were washed twice with 500 µl cold PBS. For DNA content analysis, cells were resuspended in 1 ml of 50 mM sodium citrate (pH 7.2) containing RNase A (0.2 mg/ml) and incubated overnight at 37 °C, followed by proteinase K treatment (0.4 mg/ml) for 1-2 h at 50 °C. Cells were stained with 16 µl of 1 mg/ml propidium iodide (Invitrogen) at room temperature for 30 min, briefly sonicated, and analyzed by flow cytometry. A minimum of 50,000 cells per strain was analyzed on a flow cytometer, and data were processed using FlowJo.

### dNTP pool quantification

Cells were grown to log phase in 20 ml SCD medium. One milliliter of culture was used to measure cell density with a Coulter counter, while 15 ml of cells were collected, pelleted, and snap-frozen in liquid nitrogen. We then used a methanol:chloroform extraction to measure pyrimidines. Internal standards (IS) for dATP, dGTP, dCTP and dTTP were spiked in. After extraction and drying down, the samples were resuspended and analyzed using LC-MS based on MRM (Multiple Reaction Monitoring). We used a Shimadzu UPLC Nexera II interfaced with a Sciex QTRAP 6500+ mass spectrometer equipped with a TurboIonSpray (TIS) electrospray ion source. An external standard curve was run using known concentrations of dATP, dGTP, dCTP, dTTP, and IS. The standard curves were used to calculate the concentration of the dNTP in the samples from the peak area detected.

### Sort-Seq replication timing profiling

We assessed the replication timing of the YAC-fused strains using Sort-Seq as described in^35^. Overnight cultures were diluted to inoculate 3 × 10⁷ cells into 25 ml YPAD and grown at 30 °C to a density of 0.7-1.5 × 10⁷ cells ml^-1^. Cells were collected by centrifugation (2,000 × g, 5 min, 20 °C), washed twice with distilled water, and fixed in 70% ethanol overnight at 4 °C. Fixed cells were washed twice with 50 mM sodium citrate (pH 7.2) and resuspended in 1 ml of the same buffer. RNA and protein were removed by sequential incubation with RNase A (250 µl of 10 mg ml^-1^, 1 h at 37 °C) and proteinase K (100 µl of 20 mg ml^-1^, 1 h at 55 °C). Cells were stained in 2.5 ml 50 mM sodium citrate containing 25 µl 10% sodium azide and 5 µl SYTOX^TM^ Green Nucleic Acid stain (Invitrogen) for ≥1 h at room temperature. Before sorting, samples were dispersed by two 3 s sonication pulses (20 W). Cells were then sorted on a FACSAriaIII Sorter (BD). DNA-content distributions were analyzed using a 488-nm laser with 530/30 filter. DNA fluorescence histograms were used to set the gates for sorting and isolate S- and G2-phase cells while excluding debris and doublets.

We sorted 7.5 million cells from replicating and non-replicating subpopulations. The purity of the sorted cell fractions was confirmed by flow cytometry. Sorted cells were pelleted after addition of ethanol to a final concentration of 30% (v/v). Cell pellets were resuspended in 1.2 M Sorbitol, 0.2 M Tris-HCl pH 8.0, 20 mM EDTA pH 8.0, 0.001% v/v β-mercaptoethanol and spheroplasted with 1 mg ml^-1^ Zymolyase for 30 min at 37°C. The reactions were supplemented to 0.42% w/v SDS, 85 mM NaCl, 83.3 µg ml^-1^ RNase A, and 0.33 mg ml^-1^ proteinase K and incubated for 2 h at 55°C. After incubation, samples were mixed with an equal volume of phenol:chloroform:isoamyl alcohol (25:24:1, pH 7.5-8.0, Carl Roth), followed by subsequent extractions with an equal volume of phenol:chloroform:isoamyl alcohol and chloroform. DNA was precipitated using 100% ethanol at −20 °C overnight, washed once with 70% ethanol, and eluted in 100 µl TE buffer (10 mM Tris pH 8.0, 1 mM EDTA). DNA samples were submitted for high-sensitivity library preparation and sequencing. Reads were pre-processed to remove low-quality sequences, aligned using Bowtie2 (https://github.com/benlangmead/bowtie2) and processed using a modified localMapper pipeline (https://github.com/DNAReplicationLab/localMapper/). Read counts were summarized in 1-kb genomic bins and replication profiles were generated using the Repliscope R package (https://cran.r-project.org/web/packages/Repliscope/).

### Flow cytometry to determine the yeast genomes and YAC DNA per cell

Cells were cultured in either SCD or SCGE medium to log phase (OD_600_ = 0.1-0.4). DNA content was quantified by flow cytometry to estimate the amount of DNA per cell (asynchronous cells are in a mixture of cell cycle phases). Cells (400 µl) were fixed by adding 1 ml 100% ethanol at 4 °C for ≥1 h, washed twice with 500 µl PBS, and resuspended in 1 ml of 50 mM sodium citrate (pH 7.2) containing RNase A (0.2 mg/ml). Samples were incubated at 37 °C overnight, followed by proteinase K treatment (0.4 mg/ml) at 50 °C for 1-2 h. After protein digestion, propidium iodide was added. Cells were sonicated, and DNA content was measured for >50,000 events using an Attune NxT flow cytometer. DNA (Genome copy number) per cell was calculated after gating for single cells using FlowJo, and this value was used to convert ChIP-seq occupancy normalized by spike-in controls to occupancy per cell. Then yeast genomes per cell should be:

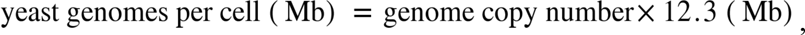

while YAC DNA per cell should be:

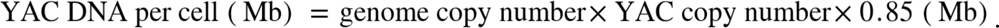

The yeast genomes per cell and YAC DNA per cell are used to calculate RNAPII occupancy and also mRNA amount (see section below).

### Mass spectrometry-based proteomic analysis

Yeast cultures (10 ml) were grown to log phase (OD_600_ ∼0.2), harvested, snap-frozen, and stored at −80 °C. Cell pellets were resuspended in lysis buffer (50 mM Tris-HCl pH 8.0, 150 mM NaCl, 0.2% Tergitol, 5 mM EDTA) supplemented with 1 mM PMSF and cOmplete protease inhibitors (1 tablet per 10 mL), and lysed using a FastPrep-24 (4 °C, 5.5 m s⁻¹, 35 s). Lysates were clarified by centrifugation (15,000 × g, 5 min, 4 °C), and the supernatant was collected. Proteins were denatured with 1% SDS and reduced with 5 mM DTT at 65 °C for 15 min, followed by alkylation with 15 mM iodoacetamide at room temperature for 15 min. Proteins were precipitated by addition of >3 volumes of precipitation solution (acetone:ethanol:acetic acid, 50:49.9:0.1), incubated on ice for 15 min, and pelleted by centrifugation (17,000 × g, 5 min, 4 °C). Pellets were washed once with precipitation solution and resuspended in 75 µL urea buffer (8 M urea, 50 mM Tris-HCl pH 8.0), followed by dilution with 225 µl 150 mM NaCl. Proteins were digested overnight at 37 °C with trypsin (1:50 enzyme-to-protein ratio) with agitation. Digestion was terminated by addition of 15 µl 10% trifluoroacetic acid (TFA), and samples were clarified by centrifugation (17,000 × g, 2 min). Peptides were desalted using C18 cartridges (Waters, WAT054955) and stored at −20 °C. Desalted peptides were analyzed on a Fusion Lumos mass spectrometer (Thermo Fisher Scientific) coupled to an EASY-nLC 1200 system. Separation was performed by capillary reverse-phase chromatography on a 25 cm column (75 µm inner diameter) packed with 1.6 µm C18 resin (Ionopticks, AUR2-25075C18A) using a 125 min stepped linear gradient at 300 nL min⁻¹ (3-27% buffer B over 105 min, 27-40% over 15 min, 40-95% over 5 min, followed by 90% buffer B for 5 min; buffer B: 0.1% (v/v) formic acid in 80% acetonitrile). The column temperature was maintained at 50 °C. Data were acquired in data-dependent acquisition mode using Xcalibur (v.4.4.16.14, Thermo Fisher Scientific) with advanced peak detection enabled. MS1 survey scans were collected in the Orbitrap mass analyzer (profile mode, 375-1,500 m/z, resolution 240,000 at m/z 200, normalized AGC target 250%, maximum injection time ‘Auto’). Precursor ions were fragmented by higher-energy collisional dissociation (normalized collision energy 31), and MS2 spectra were acquired in the ion trap (scan rate ‘Turbo’, isolation window 0.7 m/z, AGC target ‘Standard’, maximum injection time ‘Auto’). Dynamic exclusion was set to 30 s to minimize repeated sequencing, and the maximum duty cycle was 1 s.

Raw files were searched using the Andromeda engine in MaxQuant (v.2.4.2). Variable modifications included oxidation (M) and protein amino-terminal acetylation, whereas carbamidomethyl (C) was set as a fixed modification. A maximum of five modifications per peptide was allowed, and digestion was set to trypsin with proline blocking. Database searches were performed against the UniProt yeast proteome (UP000002311_559292) with a minimum peptide length of seven amino acids. A 1% false discovery rate (FDR) was estimated using a reverse decoy proteome. MaxLFQ was used to determine the summed ion intensity for each protein. MaxLFQ intensity was then used to calculate proteins concentration ratios between strains.

To determine whether the RNA transcribed from YAC DNA produced a protein product, we used the frequency with which we matched a known yeast peptide to acquired fragmentation spectra as a proxy for the presence of non-yeast (i.e., YAC-encoded) protein. Proteomics data acquired from yeast with no foreign YAC DNA have peptide-spectrum match percentage of ∼30%, and the presence of an increasing number of YAC integrations did not alter this percentage.

### Spike-in normalized RNA-seq for quantification of global mRNA amount per cell

To quantify the amount of mRNA per cell, *S. cerevisiae* cultures were grown in SCD medium (10 ml) to an OD_600_ of 0.2-0.4. Cells were collected by centrifugation (14,000 rpm, 30 s), snap-frozen in liquid nitrogen, and stored at −80 °C until processing. *C. glabrata*, used as a spike-in control, was cultured and processed under identical conditions. Pellets were thawed, resuspended in cold PBS, and mixed at a ∼2:1 ratio (*S. cerevisiae*:*C. glabrata*, by OD_600_). From each mixture, 20 µl was reserved for genomic DNA (gDNA) extraction, which was performed using the YeaStar Genomic DNA Kit (Zymo Research) according to Protocol 1. gDNA libraries were prepared commercially and sequenced on an Illumina platform (paired-end, 2 × 150 bp) to a depth of ∼1 million reads per sample. The remaining cell mixture was treated with 300 µl TRI Reagent (Zymo Research) for RNA isolation. Cells were lysed by bead beating (FastPrep 24; 4 °C, 5.5 m s⁻¹, 35 s), debris was removed (14,000 rpm, 1 min), and RNA was purified with the Direct-zol RNA Microprep kit (Zymo Research). We removed rRNA by rRNA depletion, not Poly(A) selection for mRNA. This resulted in the final RNAseq data including yeast mRNA and non-coding RNA. Libraries were sequenced on an Illumina platform (paired-end, 2 × 150 bp), generating ∼18 million reads per sample. For genomic DNA sequencing, 20 µl of the same cell mixture was processed with the YeaStar Genomic DNA Kit (Zymo Research) according to Protocol 1. Libraries were sequenced on an Illumina platform (paired-end, 2 × 150 bp) with ∼2 million reads per sample.

Through spike-in RNA-seq, we can calculate the relative mRNA amount per genome of yeast as:

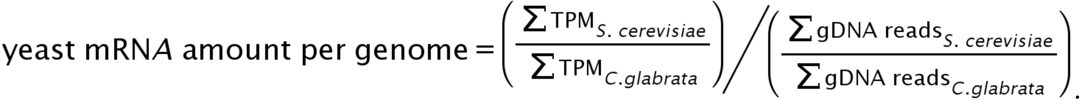

To calculate the total mRNA amount and concentration, we measured the mean cell volume by Coulter counter, and measured the genomes per cell by flow cytometry (see section above). Relative yeast mRNA amount per cell was then calculated as:

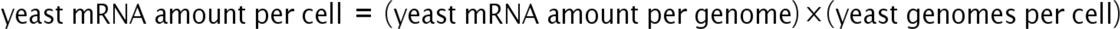

Then, the total yeast mRNA concentration was obtained by dividing mRNA amount per cell by the mean cell volume so that:

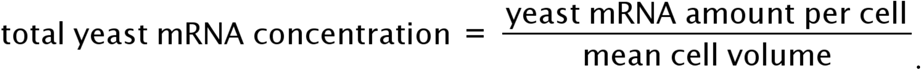

### Spike-in normalized ChIP-seq to determine RNAPII occupancy

Spike-in normalized ChIP–seq was adapted from ^28,54^. *S. cerevisiae* cultures were mixed with *C. glabrata* spike-in cultures (grown under identical conditions) at an OD_600_ ratio of 1:2 and were then fixed within 5 s by adding 1% formaldehyde for 15 min. Fixation was quenched with 0.125 M glycine for 5 min, followed by two washes in ice-cold PBS. Pellets were snap-frozen and stored at –80 °C. Thawed pellets were lysed in FA lysis buffer (50 mM HEPES-KOH pH 8.0, 1 mM EDTA, 150 mM NaCl, 0.1% sodium deoxycholate, 1% Triton X-100, 1 mM PMSF, cOmplete Protease Inhibitor) using ceramic beads in a FastPrep 24 homogenizer (MP Biomedicals). Lysates were adjusted to 1 ml and sonicated twice for 8 min each (10 s on/20 s off) using a 1/8″ microtip (Qsonica Q500) in a −20 °C ethanol bath. Debris was pelleted, and the supernatant was collected for input or ChIP. For each ChIP sample, 40 µl of Protein G Dynabeads (Invitrogen) were blocked in PBS containing 0.5% BSA for 40 min at room temperature, pre-bound with 10 µg of antibody (anti-Rpb1 clone 8wG16) in PBS for 40 min at room temperature, and then washed twice with PBS. Beads were then incubated with 0.5 ml of supernatant at 4 °C overnight. Beads were washed twice in FA lysis buffer and three times in high-salt FA lysis buffer (50 mM HEPES-KOH pH 8.0, 1 mM EDTA, 500 mM NaCl, 0.1% sodium deoxycholate, 1% Triton X-100, 1 mM PMSF), then eluted in ChIP elution buffer (50 mM Tris-HCl pH 7.5, 1% SDS, 10 mM EDTA) at 65 °C for 20 min. Simultaneously, 15 µl of input was directly combined with 115 µl of ChIP elution buffer. Crosslinks were reversed at 65 °C for 5 h, followed by RNase A (37 °C, 1 h) and proteinase K (65 °C, 2 h) treatments. DNA was purified using the ChIP DNA Clean & Concentrator kit (Zymo Research) and sequenced by paired-end (2 × 150 bp) Illumina sequencing, generating ∼18 million reads per sample.

To determine the both yeast RNAPII occupancy amounts, we need to calculate the yeast occupancy per yeast genome first, so that:

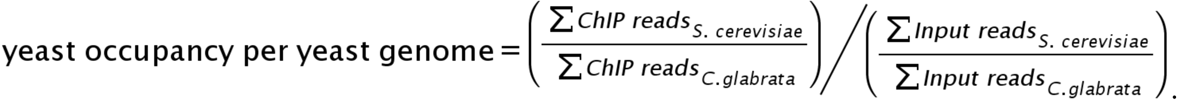

Similarly, the YAC occupancy per YAC DNA was calculated as:

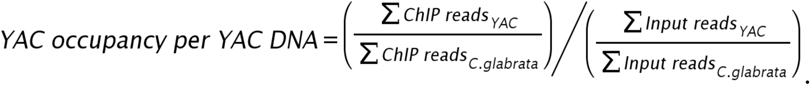

To determine both yeast and YAC RNAPII occupancy, we measured the mean cell volume by Coulter counter, and both yeast and YAC DNA per cell by flow cytometry (see section above). The relative RNAPII occupancy per cell was then obtained for the yeast genome as:

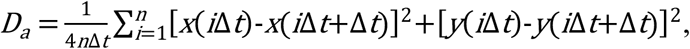

Also, the relative RNAPII YAC occupancy per cell was then calculated as:

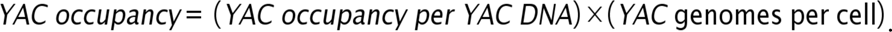

### mRNA decay measurements

Yeast strains were inoculated into fresh SCD medium lacking methionine and grown and diluted until exponential growth was reached (OD_600_ = 0.1-0.2). Transcriptional shutoff was induced by the addition of methionine to a final concentration of 1 mM. At 3-min intervals, 1.5 ml culture aliquots were collected, rapidly frozen in liquid nitrogen, and stored at −80 °C. Samples were thawed in 300 μl TRI Reagent (Zymo Research) and lysed with a FastPrep 24 instrument (MP Biomedicals; 4 °C, 5.5 m s⁻¹, 40 s). Cell debris was removed by centrifugation (13,000 rpm, 2 min) and RNA was purified using the Direct-zol RNA Microprep Kit (Zymo Research). cDNA was synthesized from 500 ng of total RNA using iScript Reverse Transcription Supermix (Bio-Rad), and quantitative PCR was performed with iTaq Universal SYBR Green Supermix (Bio-Rad). Three biological replicates were analyzed. Transcript abundances were normalized to *ACT1* and rescaled such that the initial time point was set to 1. mRNA decay constants were obtained by fitting a one-phase exponential decay model to data points collected between 6 and 18 min.

### Protein quantification by BCA assay

Cells were lysed by bead beating in lysis buffer (50 mM Tris, 5 mM EDTA, 150 mM NaCl and 0.2% Tergitol, pH 7.5, cOmplete Protease Inhibitor) on ice. Lysates were cleared by centrifugation, and protein concentrations were determined using the bicinchoninic acid (BCA) assay (Thermo Scientific) according to the manufacturer’s instructions. Briefly, samples were mixed with BCA working reagent, incubated at 37 °C for 30 min, and absorbance was measured at 562 nm using a microplate reader. Bovine serum albumin (BSA) was used to generate a standard calibration curve.

### Ribosome footprinting

Ribosome footprinting was performed following ^39^. Yeast strains were grown in 200 ml of SCD medium at 30 °C to an OD_600_ of approximately 0.5. Cycloheximide (50 µg ml^-1^) was then added to fix ribosomes for 5 min. Next, cells were harvested by centrifugation at 3,000 × g for 5 min, snap-frozen, and stored at –80 °C. Cell pellets were lysed in ice-cold buffer (20 mM Tris-HCl pH 7.4, 150 mM NaCl, 1 mM DTT, 5 mM MgCl₂, 1% Triton X-100, 25 U/ml Turbo DNase I) using a FastPrep-24 (5.5 m/s, 35 s per cycle) at 4 °C, with 5 min on-ice intervals; six cycles were performed. Lysates were clarified by centrifugation at 3,000 × g for 5 min, followed by 20,000 × g for 10 min at 4 °C. Eight hundred micrograms of RNA were diluted to 200 µl, digested with RNase I (1.5 µl, 45 min, RT), and stopped with 10 µl RNase inhibitor.

RNase-treated lysates (200 µl) were layered onto 4.5-45% sucrose gradients in polysome buffer (20 mM Tris-HCl pH 7.4, 150 mM NaCl, 5 mM MgCl₂, 1 mM DTT), centrifuged at 36,000 rpm for 2 h at 4 °C (SW41 Ti), and fractionated while monitoring A_254_. Monosome fractions were collected, treated with 1% SDS and 200 µg/ml proteinase K at 42 °C for 30 min, followed by phenol:chloroform extraction and ethanol precipitation. RNA was resuspended in Tris (10 mM, pH 8.0), denatured with RNA loading dye, separated on 15% TBE-urea gels, and the 17-34 nt region excised. RNA was eluted from gel slices in extraction buffer (300 mM sodium acetate pH 5.5, 1 mM EDTA, 0.25% SDS) overnight, precipitated, and treated with T4 polynucleotide kinase to repair 2′-3′ cyclic phosphates. Libraries were prepared using the NEBNext Small RNA Library Prep Set (E7330S) for paired-end (2×150 bp) Illumina sequencing, yielding >15 million reads per sample. Reads were trimmed with Trimmomatic v0.39 (discarding <20 nt), aligned first to *S. cerevisiae* rRNA (Bowtie v1.2, -q -v 2 -k 1 -p 4) to remove rRNA, then mapped to the combined *S. cerevisiae* (NCBI R64) and YAC (NC_060948.1) reference using Bowtie (-q -v 2 -m 1 -p 4), retaining only uniquely aligned reads for analysis.

### Single-molecule tracking to determine the active ribosome fraction

Single-molecule tracking was performed following a protocol adapted from^40^. To reduce autofluorescence, cells were cultured in low-fluorescence SCD medium for more than ten generations and collected at OD_600_ = 0.1-0.3. HaloTag labeling was performed by incubating cells with 10 nM JF-PA549 dye at 30 °C for 40 min, followed by three washes with fresh medium and resuspension in a small volume (20 µl). Cells were then applied onto SCD agarose pads prepared in gene frames on glass slides. Imaging was carried out on a Nikon N-STORM system equipped with autofocus and motorized stage control. JF549 was excited using a 561 nm laser and detected with an EMCCD camera through a high-NA oil-immersion objective under inclined illumination to suppress background. A quad-band emission filter was used, and transmitted light images were acquired for cell identification. Sparse activation of fluorophores was achieved using weak 405 nm illumination, maintaining approximately one to two detectable molecules per cell. Activated molecules were imaged using rapid 561 nm excitation (15 ms exposure), and time-lapse movies consisting of 10,000 frames were recorded with a frame interval of 37 ms. Particle detection and trajectory reconstruction were performed in ImageJ with the TrackMate plugin. Spots were identified using a Laplacian of Gaussian method (estimated diameter 1 µm). A maximum displacement of 2 µm between consecutive frames was allowed when linking trajectories, and a single-frame gap was permitted to account for transient signal loss. The diffusion coefficient per ribosome (*D_a_*) was calculated from the stepwise mean-squared displacement (MSD) of their trajectory:

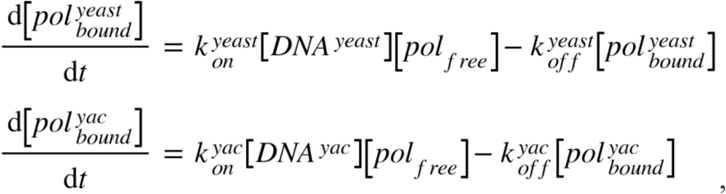

where *x(t)* and *y(t)* denote the ribosome coordinates at time *t*,Δ*t* represents the time interval between consecutive frames, and n is the number of steps in the trajectory. Trajectories with fewer than nine displacements were excluded from analysis.

### RNAPII competition model for calculating RNAPII occupancy

Our model for predicting RNAPII occupancy in either the yeast or YAC DNA is based on the mass action kinetics between free RNAPII and the yeast/YAC DNA as in^28^. The concentration of yeast DNA-bound RNAPII in the nucleus, 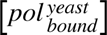, is determined by mass action kinetics of the concentration of free RNAPII, [*pol_free_*], and the concentration of yeast genome [*DNA^yeast^*]. Similarly, the concentration of YAC DNA-bound RNAPII in the nucleus, 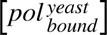, is determined by the concentration of free RNAPII and YAC DNA [*DNA^yac^*]. The dissociation of the DNA-bound RNAPII was modelled using first-order kinetics with dissociation rates 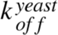 for yeast DNA-bound RNAPII and 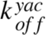 for YAC DNA-bound RNAPII so that:

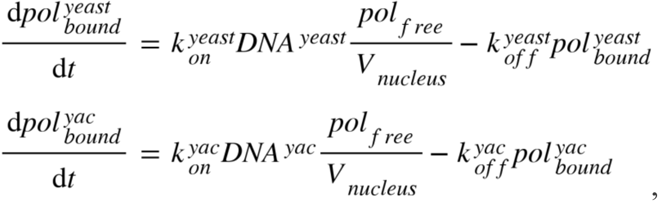

where 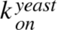 and 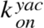 are the binding rates of free RNAPII with yeast DNA and YAC DNA respectively. The above equations can be rewritten in terms of amounts:

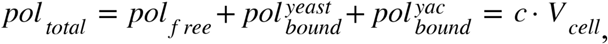

where *V_nucleus_* is the nuclear volume. Also, the total amount of RNAPII, *pol_total_*, should be the sum of free and all DNA-bound RNAPII and proportional to cell size so that:

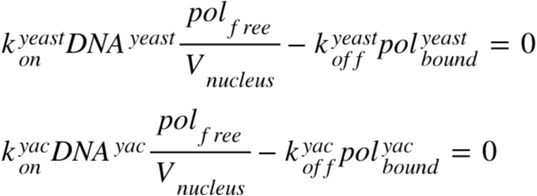

where *V_cell_* is the cell volume and *c* is the concentration of RNAPII. Since RNAPII kinetics are much faster than cell growth, their equations are at steady state so that:

The yeast DNA content and YAC DNA content can be given as:

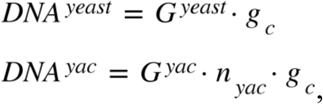

where *G^yeast^* (12.3 Mb) is the yeast genome size, *G^yac^* (0.85 Mb) is the YAC DNA size, *n_yac_* ( *n_yac_* =1, 2, …., 15.) denotes the YAC copy number, and *g_c_* denotes the yeast genome copy number. Substituting this into the above equation, the yeast DNA-bound RNAPII amount at steady state can be rewritten as:

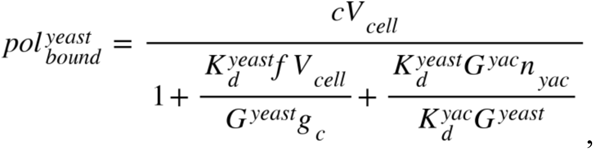

and the YAC DNA-bound RNAPII amount at steady state can be written as:

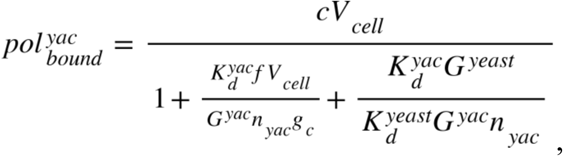

where 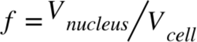 is the nuclear volume fraction, and

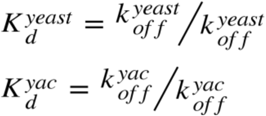

denote the equilibrium dissociation constant of RNAPII on yeast and YAC DNA, respectively. Importantly, we see that the ratio of yeast DNA-bound RNAPII and YAC DNA-bound RNAPII is:

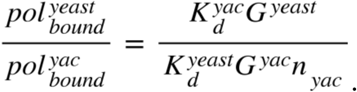

To compare our model results with measurements, we first examined how the equilibrium dissociation constant changes with YAC copy number. Based on ChIP-seq data, we found that the ratio of the YAC equilibrium dissociation constant to the yeast equilibrium dissociation constant remains consistently at approximately 1.15 across all strains, so that 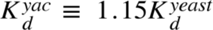. Since we independently measured the nuclear fraction, cell volume, and genome copy number, only two free parameters remain that can be fit to our data: the RNAPII equilibrium dissociation constant on yeast DNA 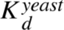, and the total RNAPII concentration *c*.

### Calculation of peptide elongation rate

The average peptide elongation rate per ribosome was estimated from the total protein synthesis rate^40^. During exponential growth, total protein mass increases proportionally to the existing mass, *m_prot_*, such that 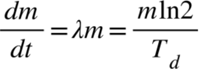, where λ is the growth rate and *T_d_* is the population doubling time. The rate of protein mass increase in a doubling time was given by the amount of ribosomes, *N_ribo_*, and the active ribosome fraction, φ*_a_*, so that the peptide elongation rate per ribosome *c_P_* is:

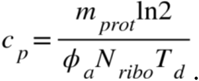

The ribosome fraction in the proteome is defined as φ*_r_*, and the average protein mass per ribosome is defined as *m_rb_*. Then the average ribosomal protein mass in the cell population is:

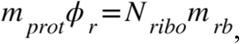

and the average elongation rate per active ribosome is given by

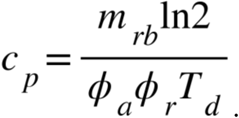

The average protein mass per ribosome *m_rb_* is ∼11,984 amino acids^55^. The remaining three parameters (ribosome fraction φ*_r_*, active ribosome fraction φ*_a_*, and doubling time φ*_d_*) were obtained from the measurements in this study.

Based on 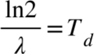, the growth rate*λ*can also be estimated as:

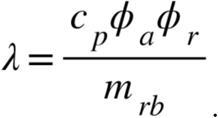

### Fluorescence in situ hybridization (FISH) to determine total mRNA concentration

Yeast cultures were grown to OD_600_ = 0.1-0.4 and fixed by adding formaldehyde (final ∼3.7%) for 45 min at room temperature. Cells were pelleted (1,600 × g, 4 min), resuspended in ice-cold fixation buffer (1.2 M sorbitol, 0.1 M K₂HPO₄, pH 7.5), and washed once before incubation with Zymolyase in the same buffer at 30 °C for 90 min to generate spheroplasts. Following two washes (400 × g, 5 min), cells were transferred to 70% ethanol and stored overnight at 2-8 °C for permeabilization. For hybridization, ethanol-treated cells were pelleted and resuspended in 100 µl hybridization solution containing 125 nM Stellaris T30-ATTO 488 probes in Stellaris Hybridization Buffer supplemented with 10% formamide, and incubated overnight at 30 °C in the dark. Cells were subsequently washed with Stellaris Wash Buffer A (including a 30 min incubation at 30 °C), followed by a brief wash in Wash Buffer B at room temperature. After the final centrifugation, cells were resuspended in 5-10 µl Vectashield mounting medium and mounted on glass slides under coverslips. Images were acquired using a Zeiss Axio Observer.Z1 wide-field epifluorescence microscope equipped with a Colibri LED illumination module and a 63×/1.4 NA oil-immersion objective. FISH signals were collected using 505 nm excitation (100% intensity, 1 s exposure), and phase-contrast images were obtained for cell boundary identification. Cell segmentation was performed using ACDC software with manual refinement. After background subtraction, mRNA concentration was estimated as total cellular fluorescence normalized by cell volume.

### Strain availability and lead contact

Yeast strains generated in this study are available from the lead contact, Jan M. Skotheim, upon reasonable request. Additional resources and reagents can also be obtained from the lead contact.

## Data availability

Sequencing data have been deposited in the NCBI Sequence Read Archive under BioProject accession PRJNA1455809. The code for modelling and simulations is available at GitHub: https://github.com/gaox0826/Competition-for-RNA-polymerase-II-between-endogenous-and-non-coding-DNA. Any other information required to reproduce the results is available from the lead contact upon request.

## Acknowledgements

We thank Matthew P. Swaffer for providing YAC strains and genome sequencing data, Jordan Xiao for guidance on live-cell microscopy, single-cell dynamics and yeast synchronization, Jacob Kim for advice on spike-in ChIP-seq and cell size measurements, and Fuying Dao for assistance in predicting replication origins in yeast YAC DNA using the iORI-PseKNC2.0 platform. We thank Amanda Amodeo, Christine Jacobs-Wagner and members of the Skotheim lab for comments on the manuscript. This work was supported by the NIH through the R35 GM134858.

## Author Contributions

N.L. designed and constructed the strain series with quantitatively increased non-coding DNA and performed most of the experiments. X.G. measured nuclear-to-cell volume ratios, performed single-molecule tracking, ribosome profiling, RNA FISH, RNAPII competition modelling, and translation modelling. M.C.L. performed mass spectrometry-based proteomic analyses. M.E.E. and G.E.N. designed and performed Sort-seq replication timing analysis. N.L., S.X., and J.M.S. conceived the study. N.L., X.G., and J.M.S. wrote the manuscript.

## Supplemental information

**Figure S1.**
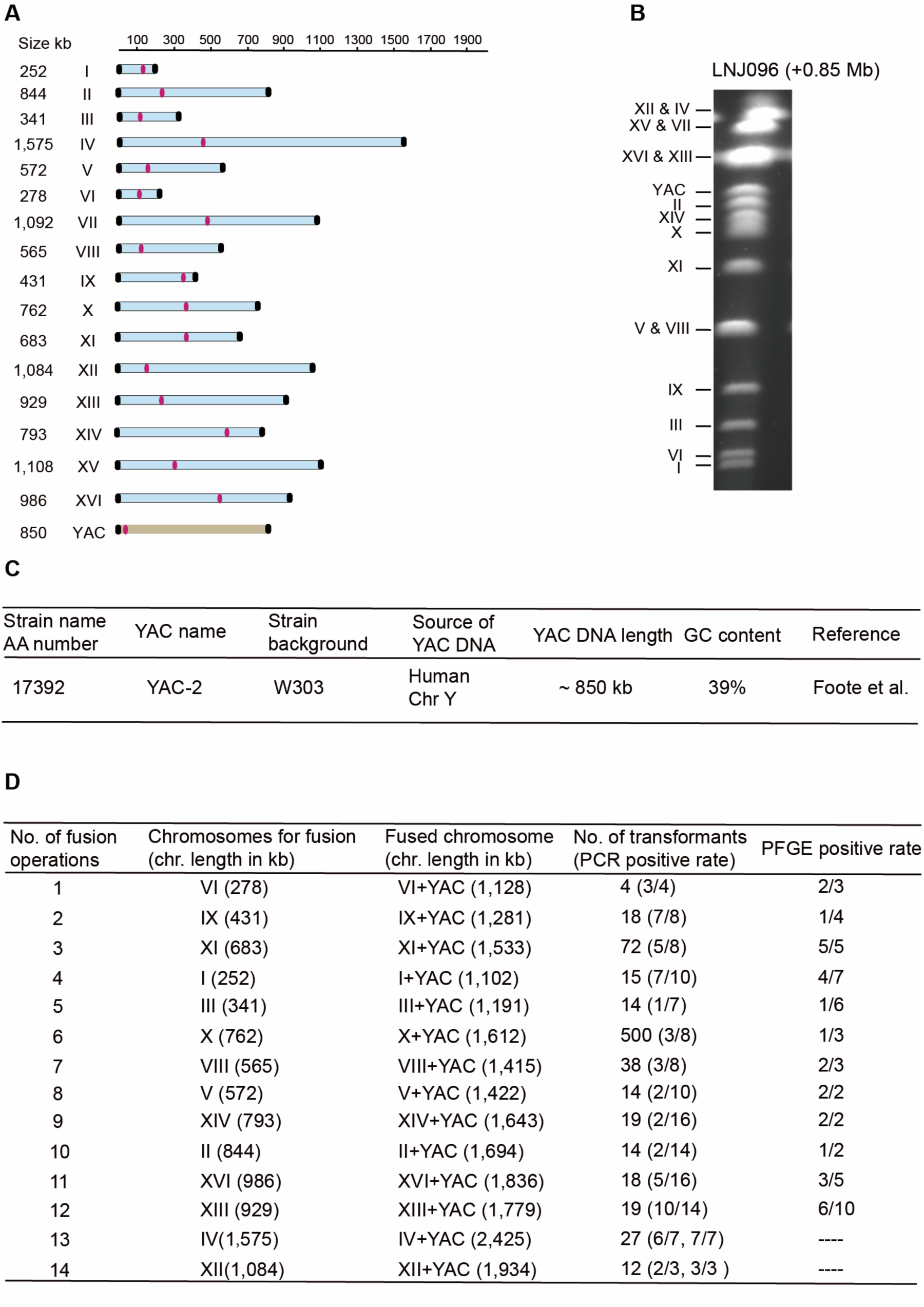
YAC origin and documentation of engineered genome expansion. **(A)** Size and position of the 16 natural chromosomes and the yeast artificial chromosome (YAC). Chromosome sizes of the W303 background strain were obtained from the genome assembly ASM216351v1, and the size of the YAC was taken from a previous study ^56^. **(B)** Chromosomes and YAC were separated by pulsed-field gel electrophoresis (PFGE). **(C)** Table of YAC strains used in this study, including source, GC content, and size. **(D)** Table of strain validation data documenting serial fusions of YACs to the endogenous chromosomes.

**Figure S2.**
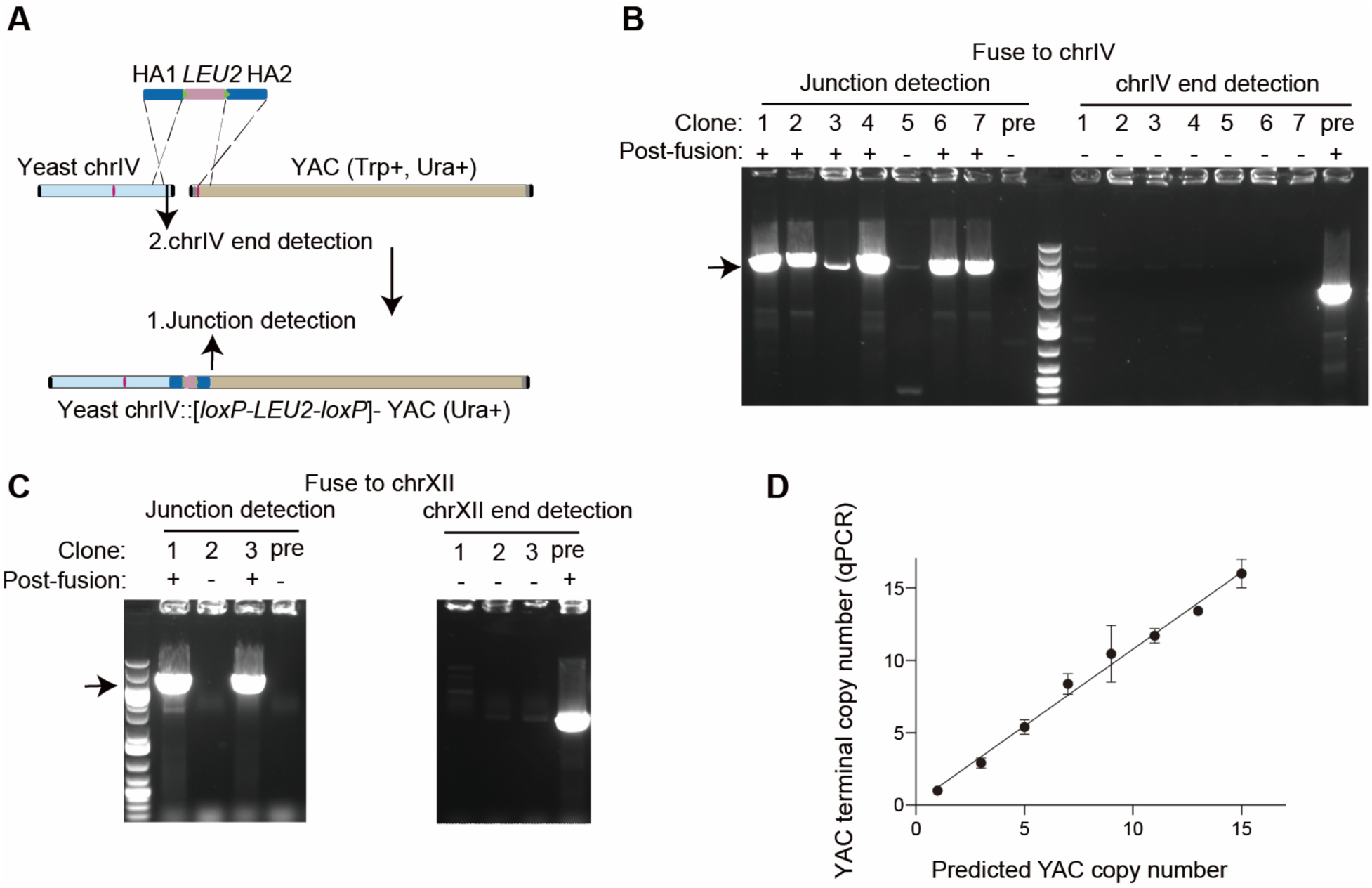
Confirmation of the gradual increase in non-coding DNA content in a series of yeast strains. **(A)** Schematic of PCR-based verification of YAC-chromosome junctions and removal of native chromosome ends. **(B)** PCR analysis of clones spanning the YAC-chromosome IV junction. Expected junction products are detected in post-fusion clones (1-7) but not in the pre-fusion control. PCR of the chromosome IV telomeric region confirms fusion by loss of the telomeric band. **(C)** PCR analysis of the YAC-chromosome XII junction, as in **(B)**, showing expected products only in post-fusion clones. **(D)** Quantification of YAC terminal copy number across strains with different amounts of non-coding DNA, measured by qPCR using *ACT1* as an internal control. Data are shown as mean ± s.d. (n = 3 technical replicates).

**Figure S3.**
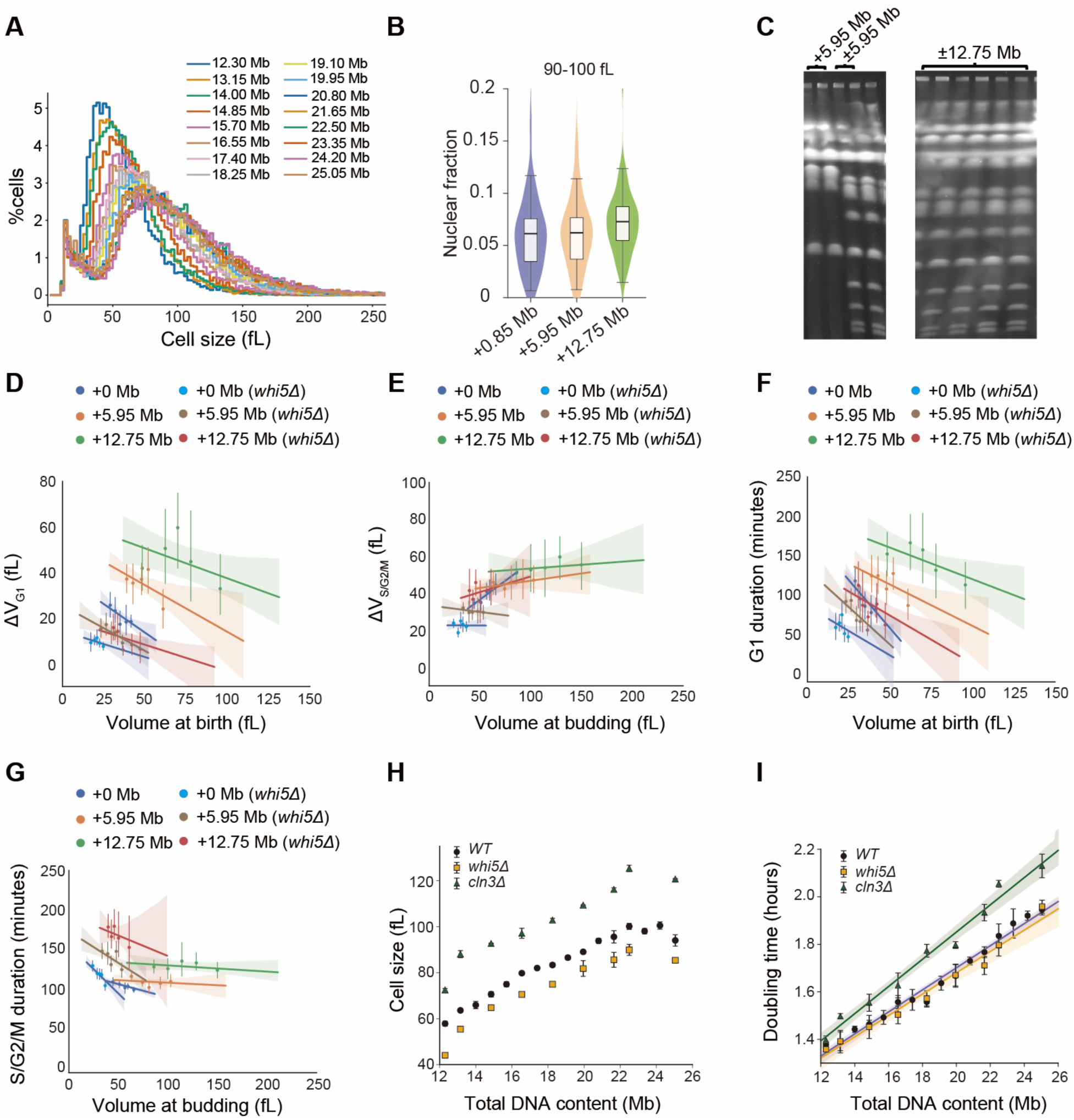
Characterization of cell size and growth in yeast strains with expanded non-coding DNA. **(A)** Representative Coulter counter profiles of cell-size distributions in strains carrying progressively more non-coding DNA. **(B)** Nuclear-to-cytoplasmic volume ratio quantified by microscopy in cells of similar size (90-100 fL) across strains with progressively increasing amounts of non-coding DNA. Boxes show the median and interquartile range, while whiskers extend to 1.5×IQR. +0.85 Mb, n = 45; +5.95 Mb, n = 24; +12.75 Mb, n = 48. **(C)** Karyotype validation of yeast strains derived from the +5.95 Mb and +12.75 Mb strains following removal of the corresponding 5.95 Mb or 12.75 Mb of human DNA. **(D)** Single-cell growth dynamics of first-generation daughters profiled in SCGE medium using microfluidics. Volume growth from birth to budding plotted against birth volume. Data are binned by equal cell numbers; dots indicate mean per bin; error bars, 95% CI; solid line, regression to unbinned data; shaded area, 95% CI. +0 Mb data is from a W303 wild-type strain^48^. +0 Mb, n = 80; +5.95 Mb, n = 104; +12.75 Mb, n = 57; +0 Mb (*whi5Δ*), n = 71; +5.95 Mb (*whi5Δ*), n = 99; +12.75 Mb (*whi5Δ*), n = 53. **(E)** Volume growth from budding to mitosis plotted against budding volume for the same cells as in **(D)**. **(F)** G1 duration plotted against birth volume for the same cells as in **(D)**. **(G)** S/G2/M duration plotted against budding volume for the same cells as in **(D)**. **(H)** Mean cell volume distributions of *whi5Δ* and *cln3Δ* mutants with increasing non-coding DNA measured by Coulter counter in SCD medium (n ≥ 2; mean ± s.d.). **(I)** Doubling times of the same mutant series with linear regression fits in SCD medium (n ≥ 2; mean ± s.d.).

**Figure S4.**
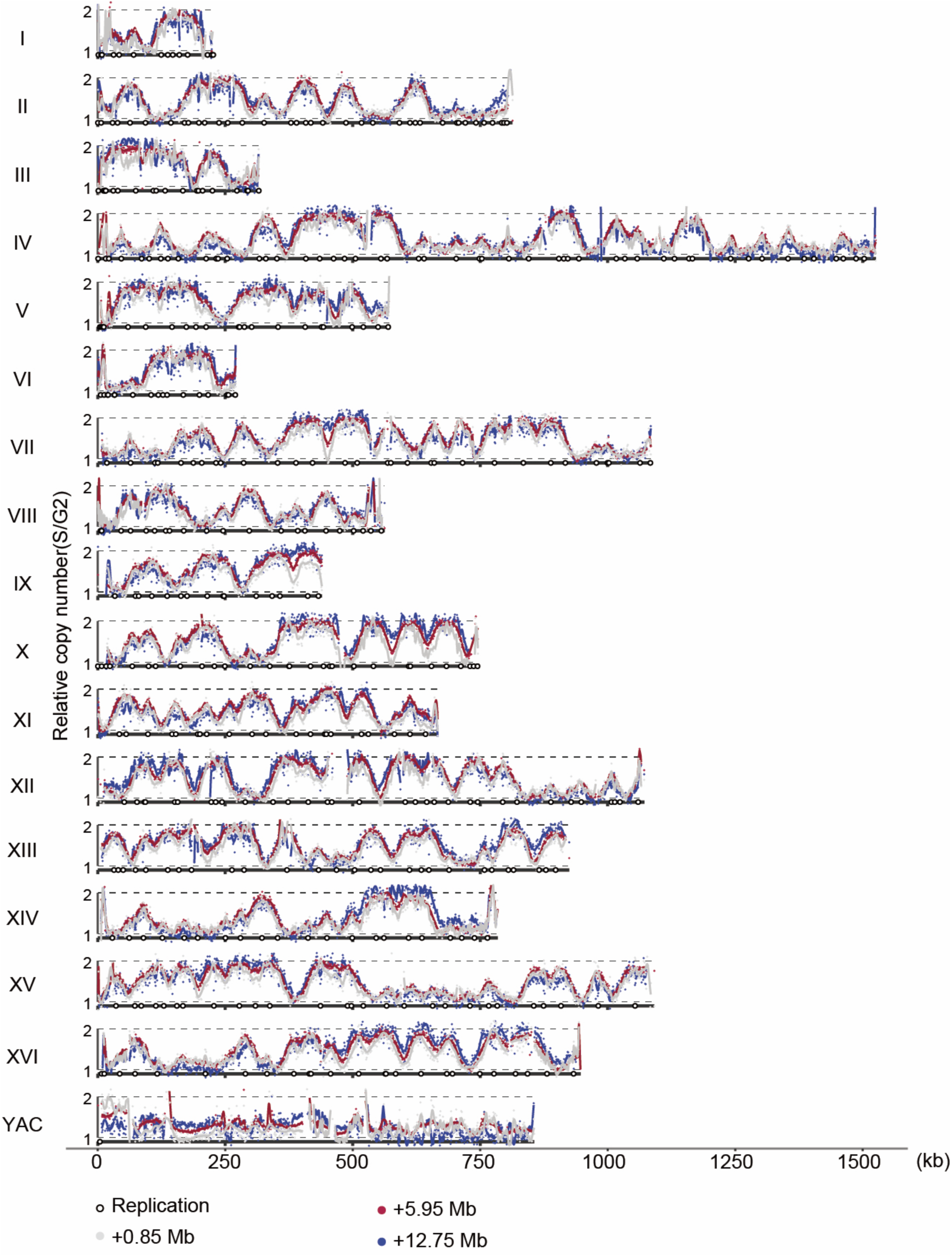
Characterization of DNA replication in yeast strains with expanded non-coding DNA. Replication profiles of yeast strains carrying +0.85 Mb (grey points), +5.95 Mb (red points), and +12.75 Mb (blue points) of additional DNA are shown together with their respective spline-smoothed fits (grey, red, and blue lines). For direct comparison, replication profiles of yeast chromosomes and the YAC were overlaid across the three strains.

**Figure S5.**
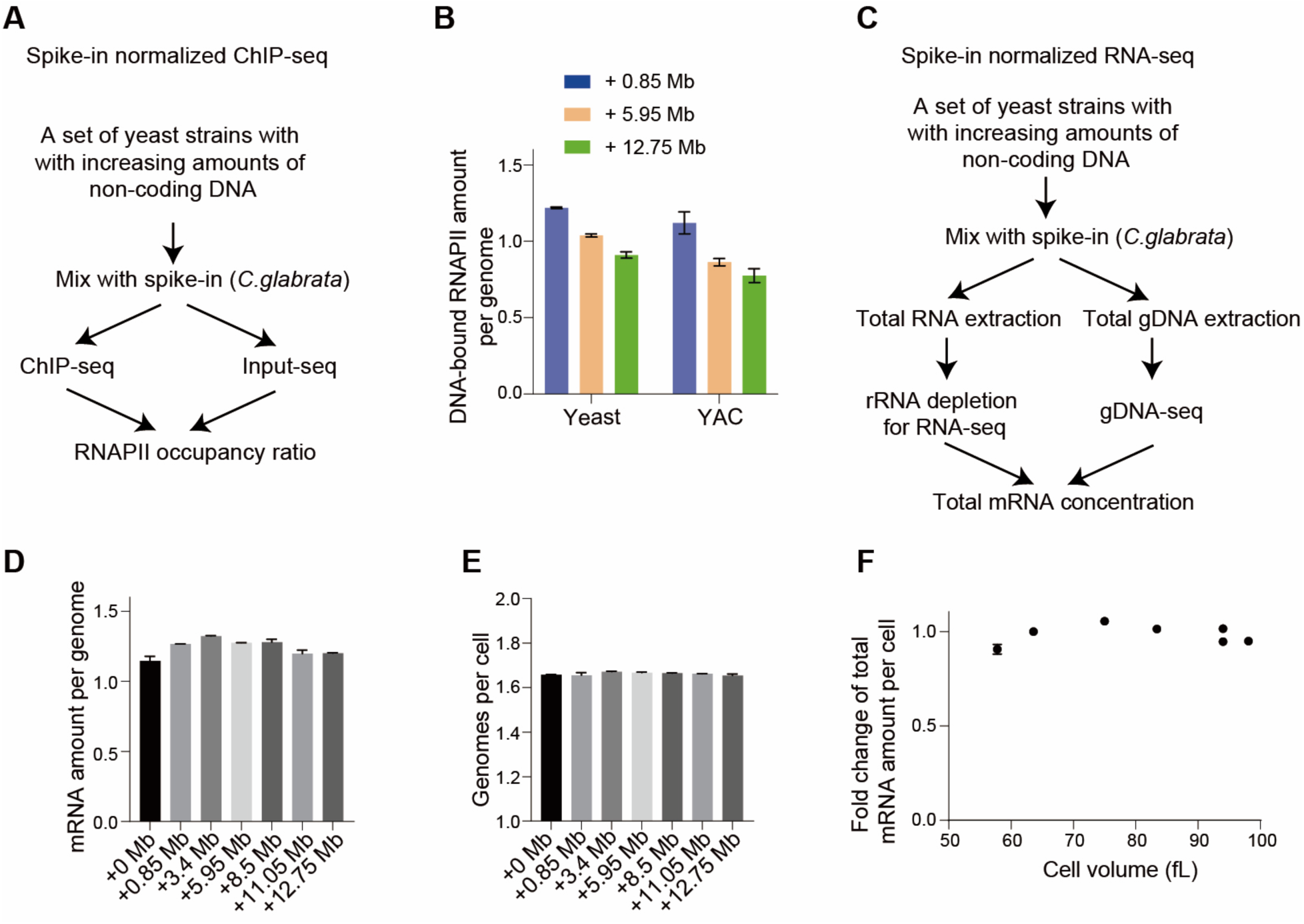
ChIP-seq and transcriptome analysis of yeast strains with expanded non-coding DNA. **(A)** Spike-in-normalized ChIP-seq workflow used to quantify global DNA-bound RNAPII. **(B)** Bar plot showing DNA-bound RNAPII per genome in strains containing varying amounts of non-coding DNA. Data are shown as mean ± range (n = 2 biological replicates). **(C)** Spike-in-normalized RNA-seq workflow used to quantify total yeast mRNA concentration. **(D)** Yeast mRNA per genome quantified by spike-in normalized RNA-seq (mean ± range, n = 2). **(E)** Average genome copy number per cell measured by flow cytometry of PI DNA-stained cells (mean ± range, n = 2). **(F)** Total mRNA amount per cell as a function of cell volume, determined by spike-in normalized RNA-seq (mean ± range, n = 2).

**Figure S6.**
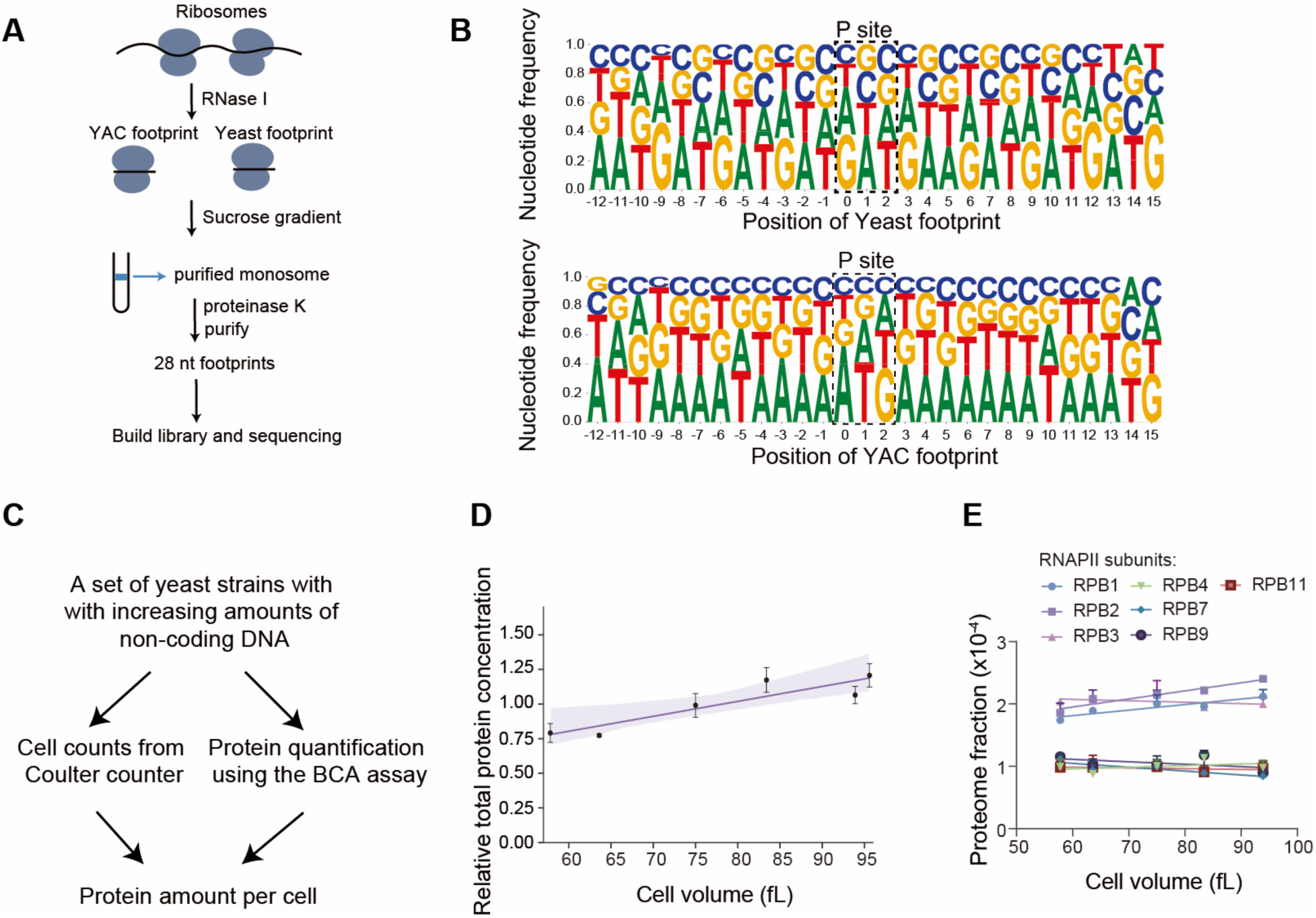
Total protein and ribosome profiling analysis of strains with increasing amounts of non-coding DNA. **(A)** Workflow schematic of ribosome footprint sequencing. **(B)** Nucleotide frequency profiles of 28-nt ribosome footprints from yeast chromosomes and YAC. The P site denotes the peptidyl-tRNA site of the ribosome, which accommodates the tRNA carrying the nascent polypeptide chain. **(C)** Schematic of protein quantification per cell by combining cell counts (Coulter counter) with total protein amounts (BCA assay). **(D)** Relative protein concentration per cell as a function of cell volume for strains with increasing amounts of non-coding DNA. (mean ± s.d., 3 biological replicates). The purple line indicates a linear regression. **(E)** Concentration of RNAPII subunits across strains of different cell sizes measured by mass spectrometry for strains with increasing amounts of non-coding DNA (n = 2 biological replicates).

**Figure S7.**
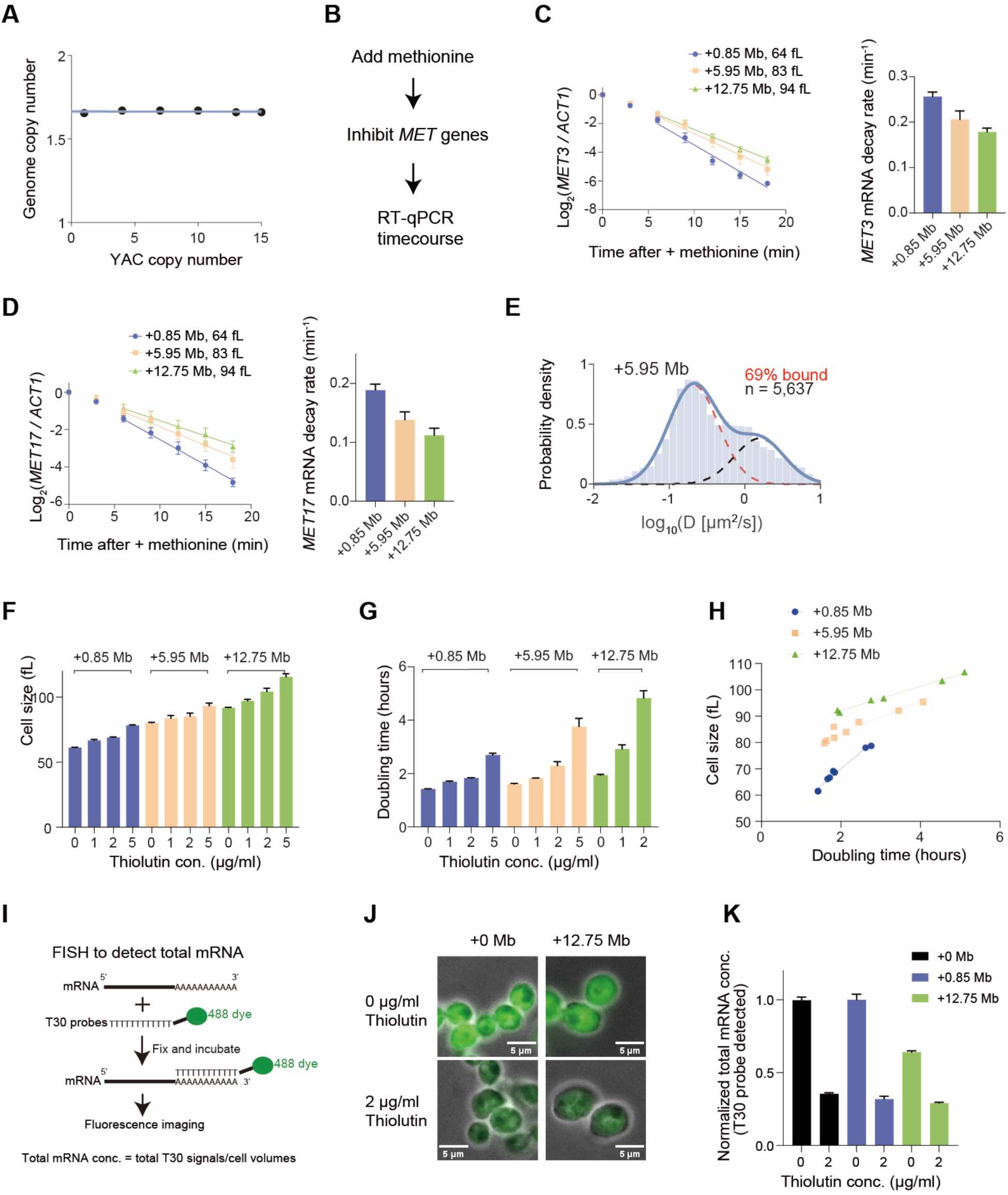
RNAPII competition model and mRNA-ribosome kinetic model. **(B)** The genome copy number in indicated strains measured by flow cytometry. Data are mean with range (n = 2). The shaded area denotes 90% CI of the fit. **(C)** Workflow for conditional transcriptional repression of *MET* genes followed by RT-qPCR analysis of *MET3* and *MET17* mRNA levels. **(D)** Left: *MET3* mRNA concentrations relative to *ACT1* mRNA concentrations after methionine addition represses *MET3* transcription in the indicated DNA-addition strains. Right: Mean (± s.e.m.) of *MET3* mRNA decay rates measured in three biological replicates. **(E)** Left: *MET17* mRNA concentrations relative to *ACT1* mRNA concentrations after methionine addition represses *MET17* transcription in the indicated DNA addition strains. Right: Mean (± s.e.m.) of *MET17* mRNA decay rates measured in three biological replicates. **(F)** The fractions of active and inactive ribosomes were distinguished using a Gaussian mixture model to fit the distributions of diffusion coefficients measured from single-molecule tracking experiments. Distribution of diffusion coefficients of ribosomes in strains with 7 YACs (n = 5,637 tracks). **(G)** Mean cell volume of asynchronously growing cultures in SCD medium supplemented with graded concentrations of thiolutin, measured using a Coulter counter (n ≥ 2; mean ± range). **(H)** Doubling times of cultures grown in SCD medium containing graded concentrations of thiolutin, calculated from OD_600_-based growth curves using linear regression (n = 2; mean ± range). **(I)** Relationship between doubling time and mean cell volume for asynchronously growing cultures in SCD medium supplemented with graded concentrations of thiolutin. **(J)** Workflow schematic of using FISH to measure total mRNA concentration. **(K)** Representative fluorescence microscopy images showing total mRNA labeled with dye in cells treated with 0 and 2 µg/ml thiolutin. **(L)** The total mRNA concentration quantified by FISH. Data are shown as mean with s.e.m. Cell numbers for each condition are as follows: +0 Mb, n = 129 (0 μg/ml thiolutin) and 347 (2 μg/ml thiolutin); +0.85 Mb, n = 305 and 341; +12.75 Mb, n = 416 and 304. Measurements at 0 μg/ml thiolutin were normalized based on mRNA concentrations determined by spike-in RNA-seq.

